# pertTF: context-aware AI modeling for genome-scale and cross-system perturbation prediction

**DOI:** 10.64898/2026.03.12.711379

**Authors:** Yangqi Su, Dingyu Liu, Vipin Menon, Bicna Song, Samuel Boccara, Nan Zhang, Huan Zhao, Jiahui Hazel Zhao, Lei Wang, Nan Hu, Mpathi Nzima, Alon Katz, Bharath Kumar Swargam, Seth A. Ament, Yarui Diao, Hanrui Zhang, Lumen Chao, Gary Hon, Danwei Huangfu, Wei Li

**Affiliations:** University of Maryland-Institute for Health Computing, North Bethesda MD 20852; Department of Pharmacology and Physiology, University of Maryland School of Medicine, Baltimore MD 21201; Developmental Biology Program, Sloan Kettering Institute; 1275 York Avenue, New York, NY 10065, USA; Louis V. Gerstner Jr. Graduate School of Biomedical Sciences, Memorial Sloan Kettering Cancer Center, 1275 York Avenue, New York, NY 10065, USA; Center for Cancer and Immunology Research, Children’s National Hospital, 111 Michigan Ave NW, Washington, DC 20010, USA; Department of Computer Science, University of Maryland College Park, College Park MD 20742; Cecil H. and Ida Green Center for Reproductive Biology Sciences, University of Texas Southwestern Medical Center, Dallas, TX, USA; The Biological Sciences Graduate Program, University of Maryland College Park, College Park MD 20742; Institute for Genome Sciences, University of Maryland School of Medicine, Baltimore MD 21201; Maryland Psychiatric Research Center, Department of Psychiatry, University of Maryland School of Medicine, Baltimore MD 21201; Department of Cell Biology, Duke University Medical Center, Durham, NC, USA; Cardiometabolic Genomics Program, Department of Medicine – Cardiology, Columbia University Irving Medical Center, New York, NY 10032; Center for Precision Medicine and Genomics Research, Children’s National Hospital. 111 Michigan Ave NW, Washington, DC 20010, USA

**Author notes:** first authors (Yangqi Su, Dingyu Liu, Vipin Menon).

## Abstract

Predicting genetic perturbation responses at a single-cell level is central to building models for cell state and disease. However, existing approaches are limited on predicting phenotypic outcomes beyond expression changes and generalizing predictions across genome-scale perturbations in biologically relevant contexts. Here we introduce pertTF, a transformer-based single-cell genetic perturbation model. pertTF was trained from a unique dataset capturing single cell expressions profiles of 30 full gene knockouts across 14 relevant cell types during human pancreatic development and beta-cell differentiation. pertTF outperforms current methods in predicting expression changes of perturbing unseen genes in unseen cellular contexts. In addition, pertTF infers perturbation-induced shifts in cell identity and population composition, an important phenotypic outcome of perturbation in many physiology and disease settings. Through transfer learning, pertTF operates in physiologically relevant systems, including primary human islets, where large-scale perturbation experiments are challenging. The generalizability of pertTF is further demonstrated by in silico pooled and single-cell CRISPR screens, capturing critical regulators of stem cells and early pancreatic cell development. These results establish pertTF as a framework for integrating large-scale single-cell perturbation data with AI models to predict genetic perturbation effects across cellular systems and disease contexts.

## Introduction

Predicting how cells respond to genetic perturbations is a fundamental challenge at the intersection of systems biology and machine learning. High-throughput and/or high-content perturbation assays (e.g., pooled CRISPR screens^1,2^, single-cell CRISPR screens like CROP-seq^3^ and Perturb-seq^4,5^) link gene perturbations to phenotypic changes and enable a systematic measurement of perturbation responses of cells. Despite these technologies, comprehensive experimental mapping of perturbation effects is still challenging or even infeasible in many cell systems, including many disease relevant cell types or systems beyond cell lines (e.g., organoids, animal models, primary cells). In addition, the combinatorial explosion of possible gene perturbations and cellular contexts far exceeds what can be assayed in the lab. These challenges motivate the development of computational models that can *in silico* predict transcriptomic responses to genetic perturbations, to prioritize experiments and guide hypothesis generation.

Measuring and predicting high-content, multimodal phenotypes to perturbations will be essential to systematically understand gene functions and their regulatory circuits. Many computational models (gene regulatory network (GRN)-focused^6,7^, deep learning^8–10^, single-cell foundation^11–16^, diffusion^17,18^, or population tracing methods^19–25^, etc.; see ref.^26,27^) focus on predicting single-cell transcriptomic responses to perturbation, thanks to the accumulating Perturb-seq or similar perturbation datasets available at the public domain. These datasets and models, however, only capture gene expression changes of perturbation within a limited gene set (e.g., transcription factors) or biological context (e.g., cancer cell lines), and struggle to capture many phenotypic outcomes beyond gene expression changes^28^. Indeed, recent benchmark studies showed that current models do not yet decisively outperform simple baselines^29,30^ and have poor generalizability in unseen cellular contexts^31^, highlighting the need to generate training datasets across contexts, improve model performance, and expand model capability to predict beyond expression changes. Importantly, cell type or composition change is one of the important phenotypes induced by genetic perturbation. Switch of cell types or states is commonly observed in many physiology and disease areas including cancer (e.g., epithelial-mesenchymal transition or EMT^32^, acinar-to-ductal metaplasia or ADM^33,34^), autism (involving shifts of glial vs neuron proportions^35^), development and tissue homeostasis. Perturbation-induced cell type/composition changes are difficult to model and predict, especially when Perturb-seq only measures perturbed cells within one or two cell types (or states).

Generalizability across different biological contexts will be critical for the application of perturbation prediction models. For example, genetically perturbing cells that are relevant to human diseases (e.g., primary cells, organoids, animal models, iPSC-derived cell types) is highly desirable. However, perturbing genes at scale in these models remains very challenging, due to multiple technical and biological factors, including a) cells may be fragile to CRISPR effectors or delivery methods (e.g., viral vectors); b) many cell populations of interest are rare; and c) many diseases do not have appropriate cellular models. As a result, most high-throughput perturbation datasets have been generated in a small set of immortalized cell lines. Methods that can transfer knowledge across cellular systems and enable in silico screens are therefore urgently needed.

Here we present pertTF (**pert**urbation **T**rans**F**ormer), a transformer-based model designed to overcome these challenges in predicting perturbation responses across different biological contexts and systems. We generated a unique training dataset that captures the single-cell transcriptome responses of fully knocking out 30 pancreas lineage regulators and diabetes risk genes. The dataset, containing over 87,000 cells, span 14 major cell types during pancreatic beta cell differentiation. PertTF builds upon the advances of this training dataset, while introducing key innovations to improve the breadth and fidelity of perturbation predictions. First, unlike models that output only gene expression profiles, pertTF is trained to predict multi-faceted cellular outcomes, including shifts in cellular identity and composition. Second, pertTF is engineered for generalization to unseen genes and contexts. By leveraging large-scale, single-cell training data and integrating biological knowledge (e.g. functional gene embeddings, known essential genes), pertTF can make plausible predictions for perturbations of unseen genes, and in cell types or disease contexts that differ from those seen during training. Through comprehensive benchmark including individual Perturb-seq, we demonstrate that pertTF outperforms many existing methods. Third, we validate pertTF in biologically relevant settings, including disease contexts such as Type 2 Diabetes. By simulating perturbations in primary pancreatic islet cells and performing in silico pooled and single-cell CRISPR screens, pertTF predictions accurately capture the functions of known β-cell regulators and match the results from pooled and single-cell CRISPR screens. Our results demonstrate that pertTF provides a comprehensive and generalizable framework for understanding genetic perturbation effects, bringing us closer to the goal of virtual perturbation screens and AI virtual cells that can guide real experimental design and biomedical discovery.

## Results

### The training dataset and pertTF model architecture

We selected human pancreas development as a model system, as it involves the specification of multiple cell types and has direct relevance to human diseases such as diabetes. To enable systematic modeling of genetic perturbation effects during human pancreatic differentiation, we generated a large, high-quality single-cell perturbation dataset (Fig. 1a), and developed a transformer-based architecture, pertTF, tailored for this task (Fig. 1b). This dataset consisted of 79 mutant hPSC clonal lines targeting 30 pancreas lineage regulators and diabetes risk genes. All clones were genotyped and pooled as a knockout village for hPSC-derived islet differentiation. Cells were collected at five differentiation stages, generating over 87,000 cells spanning 14 major cell types (see Methods).

**Figure 1.**
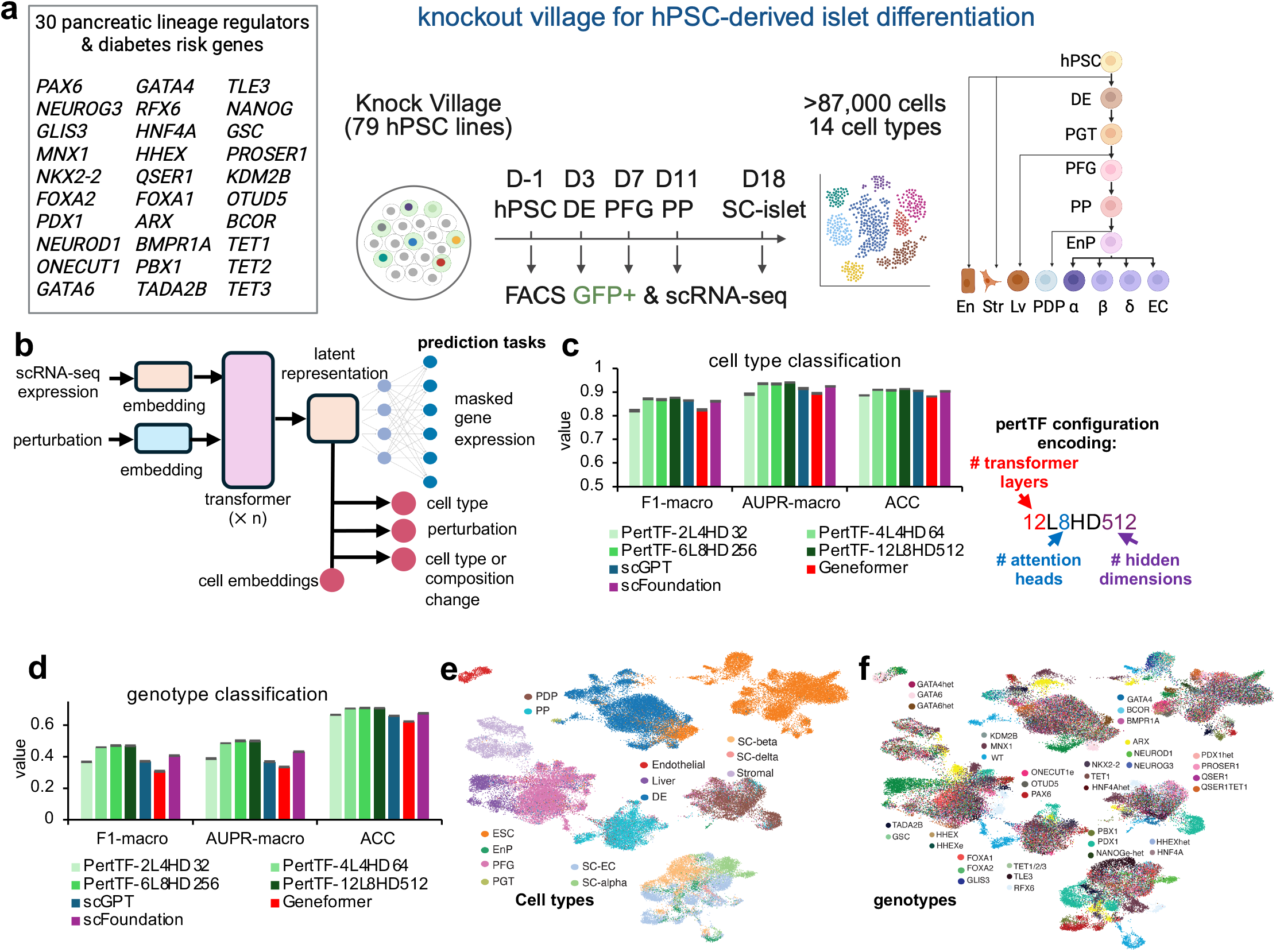
The pertTF model and training. **a.** Description of the training data generation. The entire dataset contains over 87,000 cells, covering all major cell types during pancreatic differentiation. 30 genes are knockouts and the corresponding loss-of-function genotype is confirmed via genotyping. **b**. The pertTF prediction model. pertTF takes single-cell gene expression and perturbation (optional) as input and uses different layers of transformers to generate cell embeddings, which will be further used to predict cell type, perturbation (or genotype), and cell type/composition change. In addition, masked gene expression has been incorporated into the training process. **c-d**. Performance comparison of classifying cell type (C) and perturbation (or genotype) for various pertTF settings and other foundation models, including Geneformer, scGPT, and scFoundation. Different numbers of transformer layers (L), headers (H), and dimensions (D) are tested for pertTF. For genotype classification comparison, perturbation information is masked from input. AUPR: Area Under the Precision-Recall curve; ACC: accuracy. **e-f**. The generated cell embeddings that capture cell type differences (e) and genotype differences (f). Heterozygous knockouts and enhancer knockouts are labeled as “het” and “e” in the genotype, respectively. hPSC: human pluripotent stem cell. DE: definitive endoderm. PGT: primitive gut tube. PFG: posterior foregut. PP: pancreatic progenitor. EnP: endocrine precursor. EC: enterochromaffin cells. PDP: pancreatic-duodenal progenitor. En: endothelial. Lv: liver. Str: stromal.

We designed pertTF as a multi-task transformer model that explicitly integrates single-cell gene expression profiles and perturbation information to learn a shared latent representation of cell state and genotype (Fig. 1b). The generated cell-level embeddings capture both transcriptional state and perturbation effects. During training, pertTF jointly optimizes several objectives, including masked gene expression reconstruction, cell type classification, perturbation (i.e., genotype) classification, and prediction of cell type or composition changes (Supplementary Fig. 1a-b). This multi-task formulation encourages the model to learn biologically meaningful representations that are informative for cellular context (i.e., cell type or cell state), while optimized for multiple prediction tasks. Compared with foundation models that use transformers (e.g., scGPT), pertTF introduces several key modifications in the structure to better model perturbation responses, including perturbation integration, the introduction of negative binomial (NB) log-likelihood loss and contrastive loss, adapters for phenotype/score prediction, etc. (see Methods).

The cell type and perturbation classification modules enable pertTF to accurately predict the cell type and source of perturbation, given the expression profile of a single cell. In addition, the transformer infrastructure in pertTF is compatible with many published single-cell foundation models. We tested multiple pertTF configurations with increasing model capacity (ranging from 2 transformer layers with 32 hidden dimensions to 12 layers with 512 hidden dimensions). pertTF was trained either from scratch, using only the perturbation dataset in this study for training (i.e., the 2, 4, 6 layers of transformers); or initiated using the weights of existing foundation models (i.e., the 12 layers of transformers, with initial weights copied from scGPT^15^). We compared pertTF with state-of-the-art single-cell foundation models, including scGPT^15^, Geneformer^11^, and scFoundation^16^, using held-out test cells from the perturbation dataset. Across all evaluation metrics, pertTF achieved consistently strong performance in cell type classification (Fig. 1c, Supplementary Fig. 1b). Larger pertTF models, including pertTF-12L model, exhibited improved macro-averaged F1 scores, AUPR, and accuracy, outperforming or matching fine-tuned foundation models that were pre-trained on larger external, perturbation-unaware datasets. Importantly, pertTF also demonstrated strong performance in genotype classification, a more challenging task that requires the model to distinguish perturbation-specific transcriptional signatures (Fig. 1d; perturbation information was masked from the input during training and inference).

### pertTF generates cell embeddings that jointly capture cell identity and perturbation effects

To visualize the structure of the learned latent space, we projected pertTF-derived cell embeddings using dimensionality reduction (Fig. 1e-f; Supplementary Fig. 1c-e). The embeddings formed well-separated clusters corresponding to known cell types across pancreatic differentiation, including definitive endoderm (DE), posterior foregut (PFG), pancreatic progenitors (PP), and SC-islet subtypes (Fig. 1e). In parallel, pertTF embeddings also organized cells according to perturbation genotype (Fig. 1f). Cells carrying different gene knockouts formed distinct yet partially overlapping clusters, consistent with the expectation that perturbation effects are modulated by cellular context. For example, knockouts from genotypes like *PDX1, RFX6* and *NEUROD1* lead to cells with markedly transcriptomic differences from wild-type cells in SC-islet cell types, consistent with their roles as key regulators governing different SC-islet cell types.^36^ In contrast, genotypes are not readily distinguishable via typical scRNA-seq dimensionality reduction approaches by examining top highly variable genes, where cell type differences are dominant (Supplementary Fig. 2a-b).

When perturbation information was included in the input during inference, pertTF further enhanced separation by both cell type and genotype (Supplementary Fig. 2c-g). At day 18, cells clustered cleanly by annotated cell type while simultaneously displaying structured organization by knockout identity, demonstrating that pertTF embeddings jointly encode cellular identity and genetic perturbation status within a unified latent representation. By varying the weights of loss corresponding to genotype prediction accuracy, the pertTF generated cell embedding range from completely perturbation unaware (low weight; only separating cell types), a balance between cell type and genotype (median weight), and a degenerate representation that only focus on perturbation (high weight; Supplementary Fig. 2e-g). Together, these results establish pertTF as a perturbation-aware transformer model that effectively learns biologically meaningful representations from single-cell perturbation data, outperforming existing foundation models in both cell type and genotype prediction, and laying the foundation for downstream analyses of perturbation effects across cell types and contexts.

### pertTF predicts cell type and composition changes

While most existing perturbation prediction models focus on transcriptomic changes within a fixed cellular context, a key goal of pertTF is to predict how genetic perturbations alter cell type identity and cellular composition across the developmental trajectory. To this end, we calculated the *lochNESS* score^37^, a quantitative metric that measures the enrichment or depletion of a given cell type following a genetic perturbation (Fig. 2a). Given the expression profile of an inquiry cell (usually a wild-type cell), the lochNESS score is computed from the k-nearest neighbor (k-NN) graph of cells^37,38^, by measuring the relative enrichment or depletion of neighbor cell populations (with certain gene perturbed). Positive lochNESS values indicate enrichment of a certain cell population, whereas negative values indicate depletion. The calculated lochNESS scores recapitulate known biological regulators of stem cell differentiation and cell fate decision. For example, knocking out GATA4, a critical transcription factor driving endoderm and gut tube formation, shows a depletion of PFG cells (Fig. 2b). The perturbation of HHEX leads to an enrichment of liver cell types and the corresponding depletion of PP cell types, consistent with its role as a key regulator of liver and pancreatic lineage regulator^39^ (Fig. 2b, Supplementary Fig 3a). After training, the pertTF predicted lochNESS scores match actual lochNESS scores for many genes, including PDX1, TADA2B, demonstrating strong concordance across cell types (Fig. 2c; cell types defined in Supplementary Fig. 3b), supporting the ability of pertTF to accurately model perturbation-induced compositional changes from gene expression profiles.

**Figure 2.**
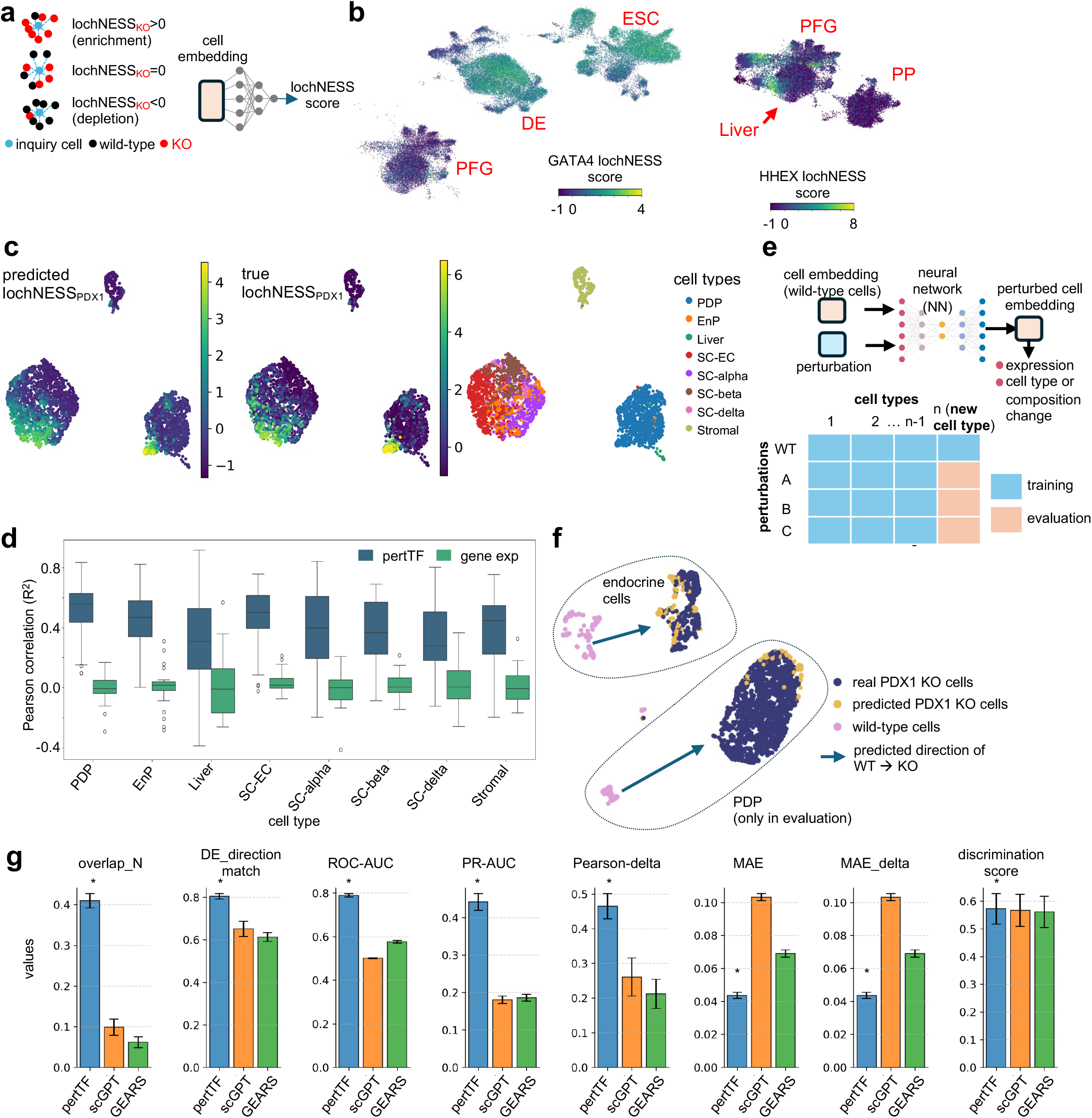
pertTF predicts cell type/composition changes and perturbations within new cell context. **a.** pertTF uses lochNESS score for predicting cell type/composition changes. **b**. The lochNESS score distribution of GATA4 and HHEX, known regulators of stem cell differentiation and liver cell fate decision. **c**. The predicted and actual PDX1 lochness scores, and the corresponding cell types in validation datasets. **d**. A comparison of lochNESS score prediction using pertTF and a naïve gene expression within different cell types. **e**. pertTF module to predict perturbed cell embeddings. Bottom: training and evaluation procedures to new cell types. **f**. Cell embeddings for predicted and real PDX1 knockout cells in various cell types, including PDP cells, whereas only wild-type cells are provided in training. **g**. Performance comparisons between pertTF and other methods including scGPT and GEARS. * indicates best performing method.

We next benchmarked pertTF against a naïve baseline approach that infers cell type changes directly from the expression of a perturbed gene, without modeling cell embeddings or perturbation dynamics. Across multiple cell types and genotypes of validation dataset, pertTF predicted scores showed substantially higher correlation with actual lochNESS score than those derived from gene expression alone (Fig. 2d; Supplementary Fig. 3c). In contrast, the gene expression–based baseline exhibited weak correlations, indicating a lower expression of perturbed gene is usually insufficient for predicting composition changes.

### PertTF outperforms existing methods in generalization to unseen cell types and perturbations

A critical challenge in perturbation modeling is generalization to new cell types (or contexts) and perturbations, a task that current algorithms are still far from sufficiently accurate^31^. Our unique perturbation datasets, spanning 14 different cell types, provide a critical training and evaluation data to address this problem. We therefore developed a pertTF module that predicts perturbed cell embeddings directly from wild-type cell embeddings and perturbation identity using a neural network trained on observed cell types (Fig. 2e). During evaluation, entire cell types (except the wild-type, unperturbed cells of that cell type) were held out from training and used exclusively for testing, ensuring that the model had no prior exposure to perturbation effects in these contexts. Using this strategy, pertTF accurately predicted the embedding shifts induced by PDX1 knockout across multiple cell types, including pancreatic-duodenal progenitor (PDP) cells that were entirely absent from the training set (except wild-type cells; Fig. 2f; Supplementary Fig. 3d). Predicted PDX1-knockout embeddings closely matched the distribution and directional shift of real knockout cells, indicating that pertTF successfully learned transferable perturbation effects that generalize beyond the training cell types.

We next compared pertTF against published deep learning methods (GEARS^8^) and foundation models (e.g., scGPT^15^, scFoundation^16^) in predicting perturbation effects in unseen cell types (Fig. 2g, Supplementary Fig. 3e) using multiple quantitative metrics, including differential expression direction matching, ROC-AUC, PR-AUC, Pearson correlation of expression changes, and mean absolute error (MAE). Across nearly all metrics, pertTF achieved the best performance across all methods (8 out of 8 metrics; Fig. 2g). In particular, pertTF showed the best discrimination of differential expression (DE) genes vs non-DE genes (measured by ROC-AUC, PR-AUC) and lower prediction error (measured by higher PCC and low MAE), highlighting its robustness in extrapolating perturbation responses to novel cellular contexts.

A central challenge in genetic perturbation modeling is predicting the effects of perturbing genes that were not included in the training dataset. To address this limitation, we extended pertTF with a module that enables inference on new gene perturbations by integrating external gene representations derived from a graph neural network (GNN), implemented in the GEARS algorithm^8^ (Fig. 3a). In this framework, perturbed genes, including those that are not observed in training dataset, are embedded using a GNN trained on Gene Ontology (GO) similarities and expression changes in a published, K562-based Perturb-seq^40^. These GNN-derived gene embeddings are then incorporated into pertTF, allowing the model to extrapolate to unseen genes. During evaluation, all cells carrying perturbations of the target gene were completely excluded from training, ensuring a strict holdout setting. We first assessed this framework by evaluating pertTF’s ability to predict the effects of PDX1 knockout, a key pancreatic transcription factor, after removing all PDX1-knockout cells from the training set. Across multiple cell types, including SC-alpha, SC-beta, PDP, and SC-EC cells, pertTF accurately predicted the perturbed cell embeddings corresponding to PDX1 loss (Fig. 3b; Supplementary Fig. 4a). Predicted PDX1-knockout embeddings closely aligned with real knockout cells and followed a consistent directional shift from wild-type to perturbed states in embedding space, demonstrating that pertTF can recover perturbation effects for a biologically critical gene without having observed any direct training examples.

**Figure 3.**
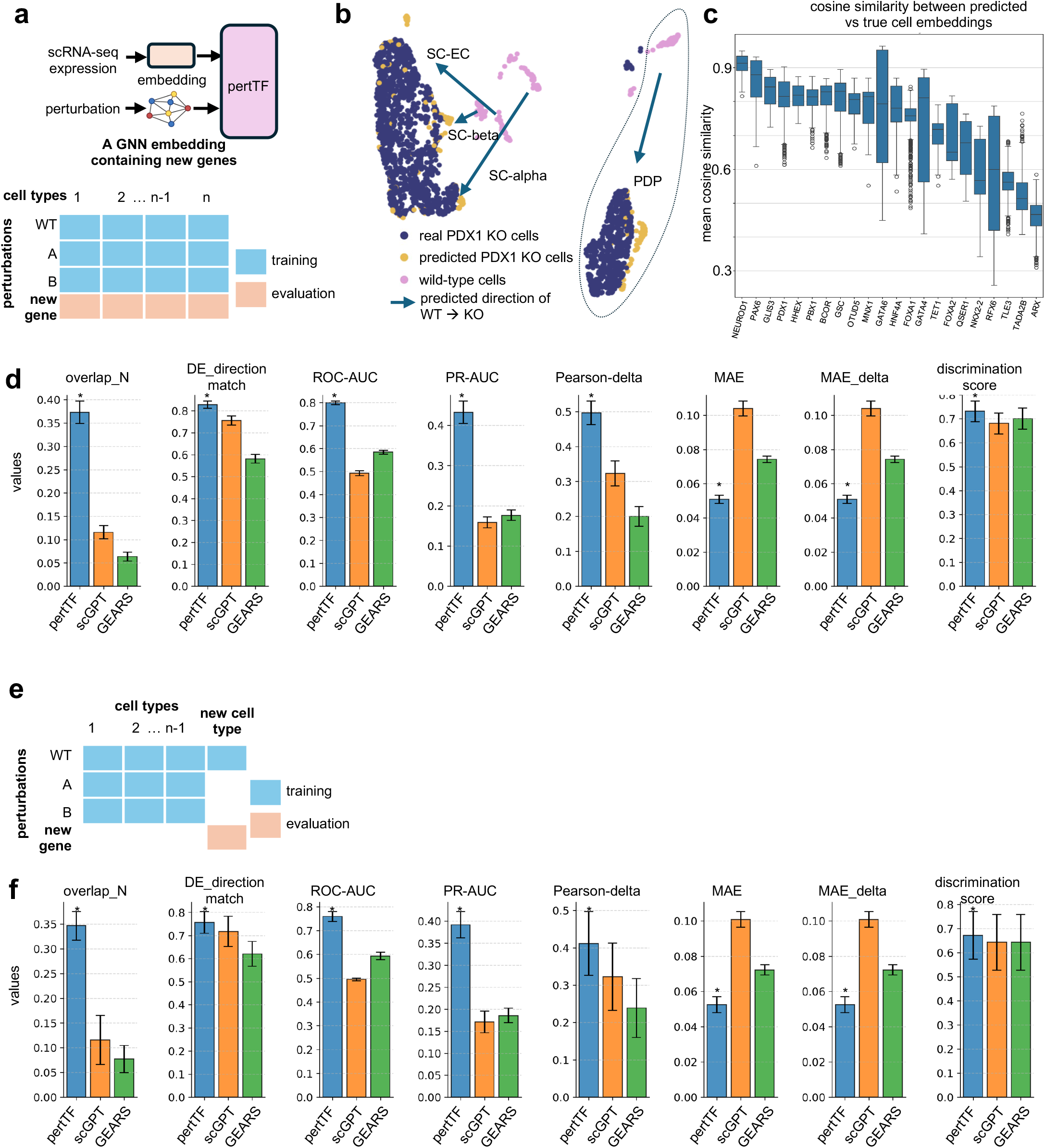
pertTF in predicting new gene perturbations. **a.** pertTF module to predict new gene perturbations using a graph neural network (GNN) embedding containing unseen genes. Bottom: training and evaluation procedures to unseen genes. **b**. Cell embeddings for predicting PDX1 knockouts across different cell types. All PDX1-KO cells are completely removed from training. **c**. A comparison of predicted vs true cell embeddings across different genotypes. **d**. A systematic evaluation of pertTF, scGPT and GEARS across different evaluation metrics in predicting gene expressions. **e**. Training and evaluation procedures for comparing unseen gene perturbations in unseen cell types. **f**. Performance comparisons of different methods.

For a systematic comparison, we performed this “leave-one-genotype-out” experiment for all 30 genotypes and compared predicted and true cell embeddings using cosine similarity (Fig. 3c). Across held-out genes, pertTF achieved higher similarity between predicted and observed embeddings than scGPT, indicating that the model captured the global structure of perturbation-induced state changes rather than overfitting to gene-specific training examples (Fig. 3c, Supplementary Fig. 4b). We next performed a systematic benchmarking of pertTF against scGPT, scFoundation and GEARS for predicting gene expression changes resulting from unseen gene perturbations (Fig. 3d, Supplementary Fig. 4c). Across multiple evaluation metrics, pertTF consistently outperformed both comparison methods in 8 out of 8 metrics (Fig. 3d), highlighting its accuracy in extrapolating perturbation effects beyond the training gene set.

We further evaluated model performance under an even more stringent setting, in which both the perturbed gene and the cell type were absent from the training data (Fig. 3e, Supplementary Fig. 4d). In this scenario, all methods were required to simultaneously generalize across genetic and cellular dimensions. Despite this increased difficulty, pertTF maintained better performance (8 out of 8 metrics) across all evaluated metrics relative to scGPT and GEARS (Fig. 3f). Notably, pertTF maintained a strong advantage over second-best performing methods, indicating that its learned representations are transferable across both genes and cell types. Together, these results demonstrate that pertTF, augmented with graph-based gene embeddings, enables accurate prediction of perturbation effects for previously unseen genes, even in novel cellular contexts. This capability substantially extends the applicability of perturbation modeling beyond experimentally assayed gene sets and represents a key step toward comprehensive in silico genetic screening.

### Independent experimental validation of pertTF predictions via CRISPRi-based Perturb-seq

We performed independent experimental validation to rigorously assess the predictive accuracy and generalizability of pertTF. We performed a CRISPR interference (CRISPRi)-based Perturb-seq experiment in induced pluripotent stem cells (iPSCs), to evaluate (1) the generalizability of our model across different perturbation strategies (full gene KO vs CRISPRi knockdown); and (2) model performance in predicting unseen genes, of which only two out of 50 perturbed genes in Perturb-seq were present in the training dataset. We designed a library containing over 400 gRNAs and performed Perturb-seq experiment. We obtained over 100,000 high-quality cells after filtering. Notably, CRISPRi induces variable perturbation efficiencies for most genes, as are revealed by clustering (Fig. 4a; Supplementary Fig. 5a) and the results from Mixscape^41^ and Perturbation-response Score^42^ (PS; Supplementary Fig. 5b-c). We therefore limited our comparison to cells with strong predicted knockdown effects (e.g., CTNNB1; Fig. 4a-b).

**Figure 4.**
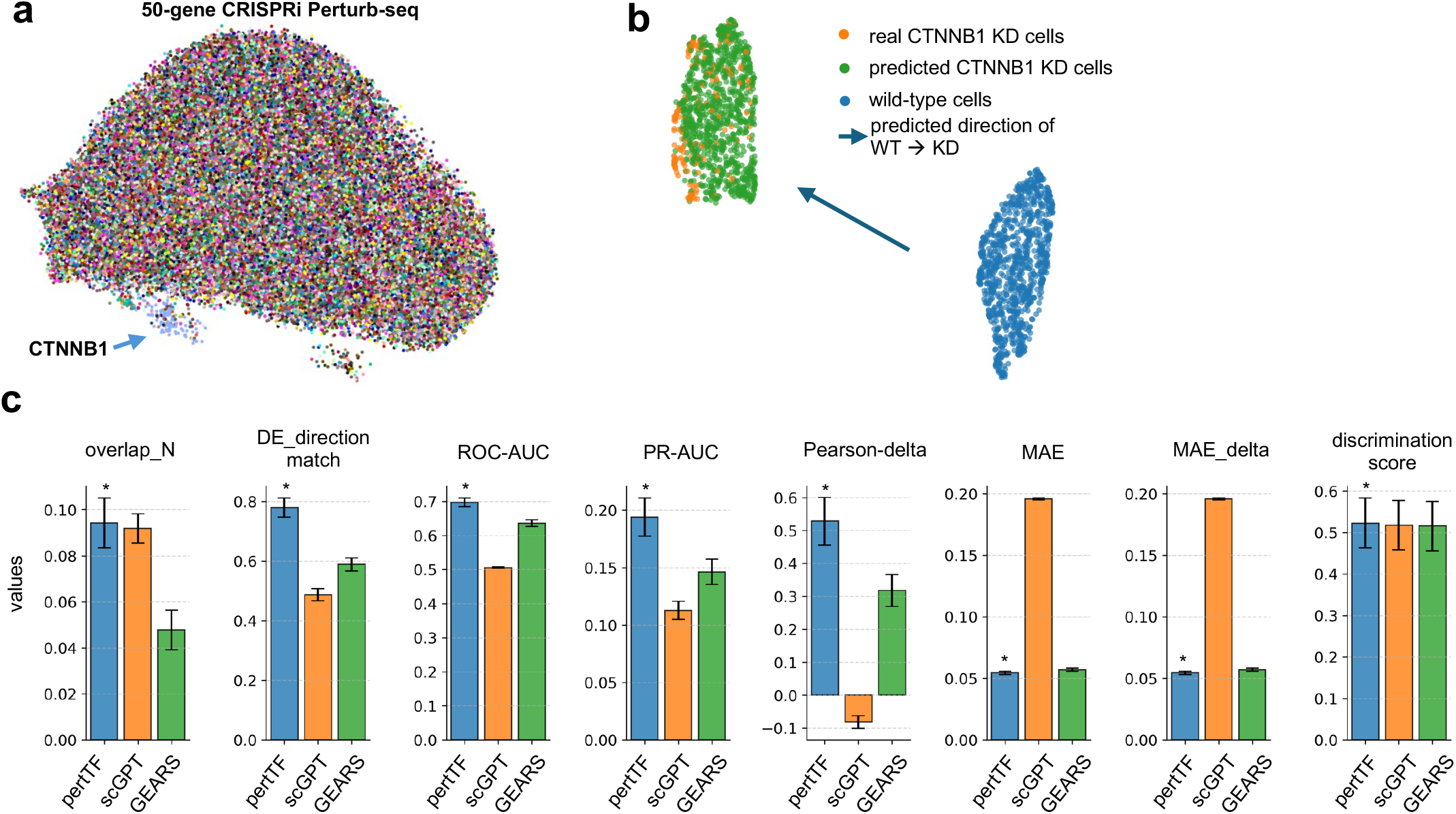
Independent experimental validations of pertTF predictions using CRISPR interference (CRISPRi)-based Perturb-seq. **a.** A UMAP presentation of all cells in the Perturb-seq, colored by perturbed genes. **b**. A systematic evaluation of expression prediction on 10 genes with strong perturbation effect (measured by Mixscape). **c**. Predicted and actual cell embeddings of CTNNB1 perturbation.

In terms of performance comparison, pertTF achieved better performance in 8 out of 8 metrics (Fig. 4c; Supplementary Fig. 5d), demonstrating that pertTF model can generalize to perturbations using different strategies and across many unseen genes. However, despite the strong performance of pertTF, all benchmarked methods performed substantially worse across all metrics than in the earlier benchmarks (Figs. 2g and 3f), indicating that predicting perturbation effects for unseen genes across different perturbation technologies is a considerably more challenging task. Notably, pertTF correctly predicted the cell embedding shift of CTNNB1, a gene where many cells show the strongest expression changes (Fig. 4b; Supplementary Fig. 5e-f). Collectively, benchmark results from both independent validations demonstrated pertTF as a stable framework that generalizes to perturbing unseen genes using different perturbation technologies.

### Transfer-learning perturbation predictions in primary human islet single-cell datasets

pertTF is trained from data generated by hPSC-derived cell types. To evaluate whether pertTF can generalize beyond cell lines to physiologically relevant human tissues or models, we applied the model to primary human islets, where genetic perturbation at scale is challenging^43^. This dataset contains scMultiome profiles of multiple non-diabetic, pre–Type 2 Diabetes (pre-T2D), and T2D donors^44^. In this setting, perturbation identities are unknown, posing a challenging inference task that requires pertTF to leverage learned perturbation signatures without explicit genetic labels. We adopted a transfer learning strategy in which a pertTF model pre-trained on pooled hPSC-derived perturbation data was fine-tuned using a small number of labeled primary islet cells (Fig. 5a). Because cell type labels are largely conserved between cell lines and primary islets, we fine-tuned our pertTF model with two objectives: predicting masked gene expression and classifying the cell type labels. Weights corresponding to other tasks (like perturbation prediction) remain frozen during fine tuning. Without fine-tuning, the pre-trained model partially captured islet cell structure but showed notable mismatches between predicted and true cell types (Fig. 5b). Fine-tuning rapidly improved alignment between predicted and true cell types, with substantial gains observed after only a few epochs and further refinement at later epochs. Quantitatively, cell type classification performance improved monotonically with both the number of primary cells used for fine-tuning and the number of training epochs (Fig. 5c). Remarkably, near-optimal precision, recall, and F1 scores were achieved using as few as 100–300 primary cells, demonstrating that pertTF can be efficiently adapted to new tissue contexts with minimal labeled data.

**Figure 5.**
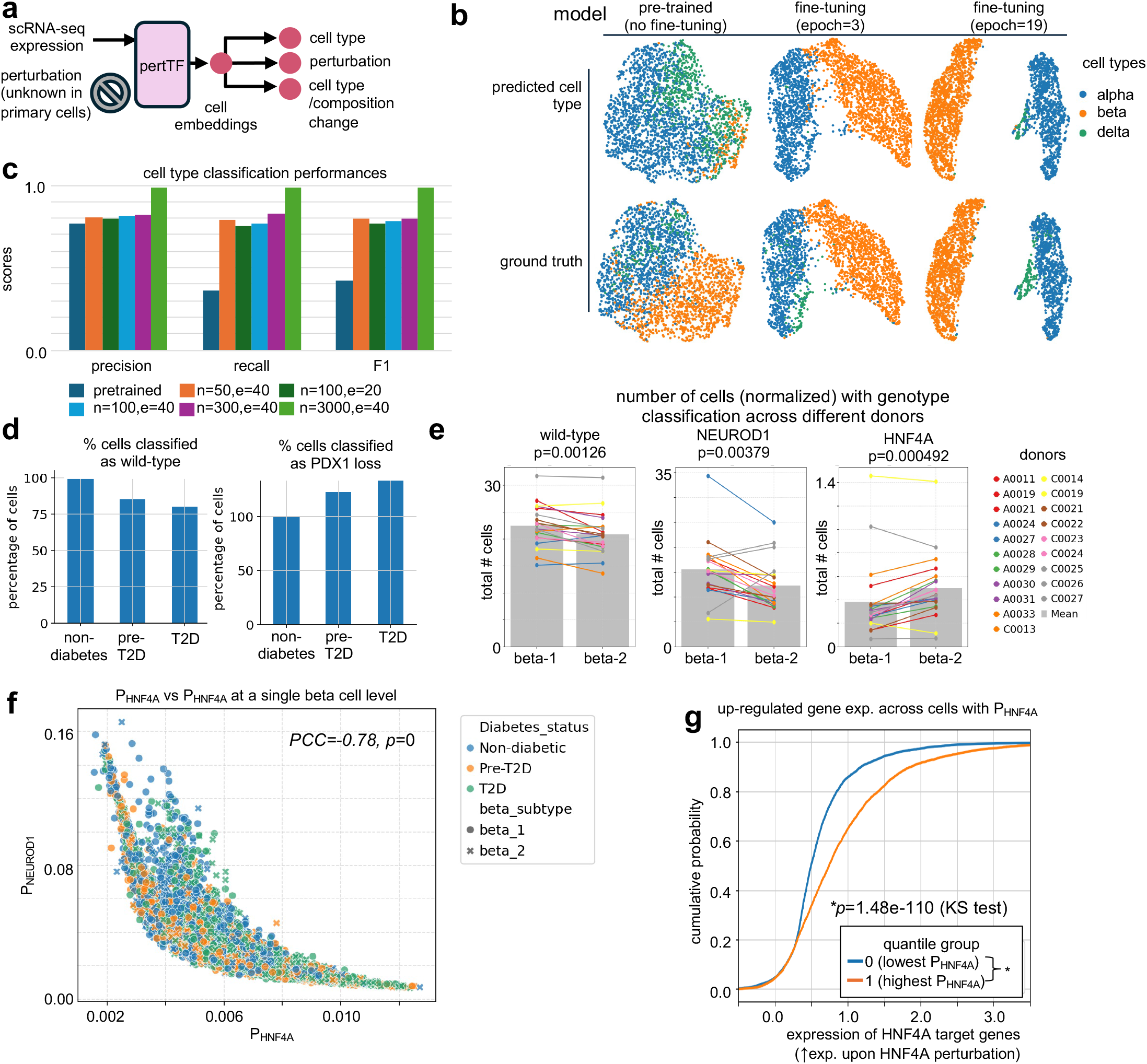
The application of pertTF to primary human islet single-cell RNA-seq datasets. **a.** Fine-tuning strategy. **b**. The cell embeddings and predicted vs true cell types for pre-trained model (no fine tuning) and different epochs of fine tuning. **c**. Performances of cell type classification using different number of primary cells (n) and different epochs. **d**. The percentage of cells classified as wild-type or PDX1 loss across different individuals with non-diabetes, pre-T2D and T2D. Cells are normalized by the numbers in non-diabetes individuals. **e**. The total number of cells classified with wild-type, NEUROD1 or HNF4A perturbations across two beta cell types within individual donors. **f**. NEUROD1 signature gene expression across cells with different groups of probabilities that have NEUROD1 perturbation (P_NEUROD1_). **g**. The fraction of cells that are predicted as RFX6 perturbation for different cell types. Cells are grouped by cell type and whether cells received RFX6 knockdown (KD) using siRNA. **h**. Cell embeddings for the predicted and real RFX6 perturbations across different cell types.

We next examined whether pertTF could infer latent perturbation states in primary islet cells. Using the fine-tuned model, we classified cells as wild-type or PDX1 loss-of-function based on predicted perturbation probabilities. Across donors, the proportion of cells classified as PDX1 loss increased from non-diabetic to pre-T2D and T2D individuals, while the proportion classified as wild-type decreased correspondingly (Fig. 5d). These trends are consistent with the known role of PDX1 dysfunction in beta-cell failure and T2D progression. Similarly, for many other known disease risk genes, the proposition of cells classified as perturbed increased in pre-T2D or T2D vs non-diabetes (e.g., GATA4; Supplementary Fig. 6a), supporting the biological relevance of pertTF-inferred perturbation states.

Beta cells are found to contain two subtypes (beta-1 and beta-2); they are driven by the expression of different set of transcription factors, whose identity were derived from the expression and motif analysis of the scMultiome data^44^. Beta-1 cells were predominant in non-diabetic donors and are driven by transcription factors like HNF1A/HNF4A/HNF4G. In contrast, beta-2 cells were more abundant in T2D donors and are driven by TCF4/NEUROD1/NFIA. Applying our model to two beta subtypes, we quantified the number of cells classified as wild-type, NEUROD1-perturbed, or HNF4A-perturbed across two beta cell subtypes (beta-1 and beta-2) within individual donors. Significant shifts in the distribution of predicted perturbation states were observed across donors for both NEUROD1 and HNF4A (Fig. 5e), indicating inter-individual heterogeneity in inferred regulatory disruption. Overall, cells classified as wild-type (unperturbed) decreased in beta-2 compared with beta-1, consistent with the identification of beta-2 as a more disease-prone beta cell type^44^. Beta-2 cells tend to have fewer NEUROD1 perturbed cells and are more likely to be perturbed by HNF4A, consistent with the key roles of NEUROD1, but not HNF4A, in beta-2 cells (Fig. 5e). Importantly, the probabilities of either NEUROD1 or HNF4A being perturbed (*i*.*e*., P_NEUROD1_ and P_HNF4A_) are strongly negatively correlated at a single-cell level (Fig. 5f), implying a mutually exclusive patterns of both beta-1 and beta-2 cell types. These results highlight pertTF’s ability to capture donor-specific perturbation landscapes within defined cell populations.

To further assess whether predicted perturbation probabilities correspond to expected molecular phenotypes, we stratified cells by their predicted probability of HNF4A perturbation (P_HNF4A_) and examined HNF4A target genes, identified from differential expression analysis in training dataset (Supplementary Fig. 6b). These gene expressions are higher in HNF4A KO cells in the training dataset, serving as stable indicators of HNF4A perturbation (Supplementary Fig. 6b), instead of a single HNF4A expression^42^. In primary cells, those with higher P_HNF4A_ showed stronger expression of HNF4A target genes, with clear separation between probability quantiles (Fig. 5g; KS test p = 1.48 × 10^-110^). Similar trends have been identified in two other key transcription factors, NEUROD1 and RFX6 (Supplementary Fig. 6c). This strong concordance between predicted perturbation likelihood and transcriptional readouts provides independent validation that pertTF predictions reflect regulatory states.

### pertTF enables in silico genetic screens

A major application of predictive perturbation models is the ability to perform *in silico* genetic screens including pooled and single-cell screens, prioritizing candidate genes whose perturbation is expected to produce a desired cellular phenotype. Using pertTF, we developed two complementary screening strategies (Fig. 6a). In Method 1, a shift of cell embeddings is predicted for perturbing each gene, and gene perturbations are ranked by the cosine similarity between predicted cell embeddings and a target perturbed population (e.g., PDX1^−^ cells). In Method 2, genes are ranked using the predicted *lochNESS* score, which directly estimates the enrichment or depletion of a target cell population upon perturbation. Both strategies provide complement approaches for in silico screens. For example, cell embedding prediction (Method 1) enables multiple downstream tasks (e.g., expression prediction), and can be integrated with many distribution alignment algorithms (e.g., optimal transport^19–24^, flow matching^25^). In contrast, directly predicting *lochNESS* value (Method 2) does not require the source/target cell population, and the *lochNESS* score provides a simple yet efficient measurement for modeling enrichments or depletions of cell populations.

We first validated the *in silico* screening framework using posterior foregut (PFG), a cell population defined by PDX1 expression^39^. Using wild-type PFG cells as the source population and experimentally observed PDX1-knockout cells as the target population, pertTF predicted perturbation effects for ∼9,800 genes and ranked them by embedding similarity (Fig. 6b). These rankings were compared against results from a published pooled CRISPR screen conducted under comparable conditions, in which cells were sorted based on PDX1-GFP expression^39^ (PDX1/GFP+ and PDX1/GFP-). pertTF successfully ranked PDX1 as the top hit and prioritized multiple known regulators of pancreatic progenitor identity, including GATA6 and MAPK1, among the top-ranked genes (Fig. 6c). This demonstrates that pertTF can recover biologically established regulators without explicit supervision from pooled screening data. Quantitatively, pertTF-derived rankings showed substantially improved agreement with pooled CRISPR screen results compared to rankings based solely on differential gene expression. Receiver operating characteristic (ROC) analysis showed that pertTF achieved a higher area under the curve (AUC = 0.79) than expression-only ranking (AUC = 0.66; Fig. 6d). Similarly, a higher area under the precision-recall curve (AUPR) for pertTF indicate improved sensitivity and specificity in identifying functional regulators of PDX1 expression (Supplementary Fig. 7a).

**Figure 6.**
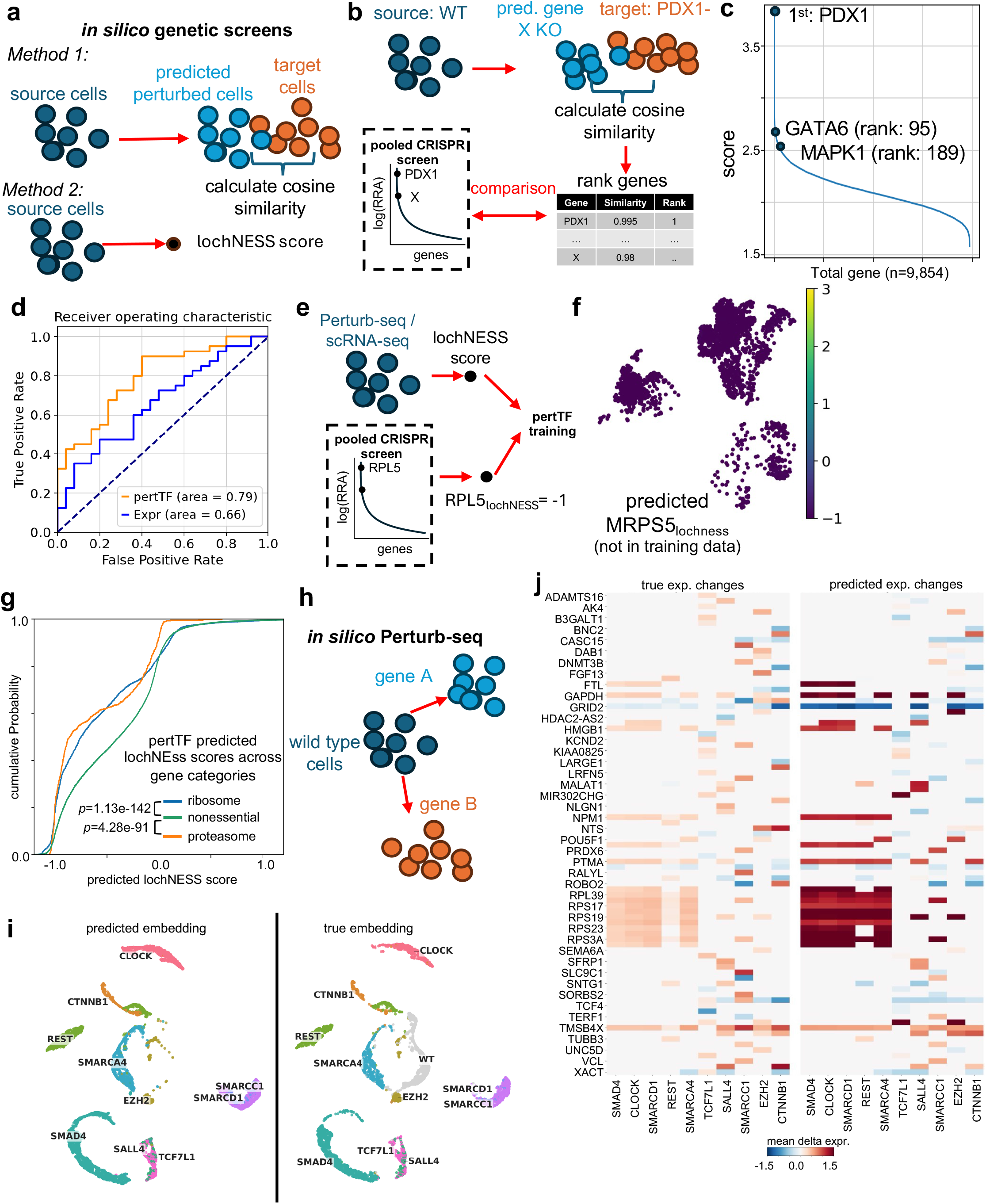
pertTF enables in silico genetic screens. **a.** Two different approaches of performing in silico screens using cosine similarity (method 1) or using predicted lochNESS score (method 2). **b**. Validation of pertTF in silico screening pipeline for critical factors of pancreatic progenitors (PP), which expresses PDX1. pertTF ranks genes whose perturbation is most similar with PDX1-cells. A published, pooled CRISPR screen of similar settings (sorting for PDX1-cells) was used as validation. **c**. Ranking of genes that most resemble PDX1-cells in PP. Other known regulators for PDX1 expression, including GATA6 and MAPK1, are marked. **d**. The ROC curve of pertTF predicted gene ranking vs. using only gene expression to generate ranking. **e**. Training strategy for pertTF in silico screens using lochNESS score, by incorporating known gene functions (e.g., known essential genes). **f**. The predicted lochNESS scores of MRPS5 which is not in training data. **g**. The cumulative distribution of lochNESS scores between non-essential genes and known essential genes (e.g., ribosomal genes, proteasomes). **h**. in silico Perturb-seq. **i**. The predicted vs true cell embeddings in Perturb-seq on genes with the strongest perturbation effects. **j**. The predicted vs true gene expression changes across the top differential expressed genes for these genes. Only genes with strongest perturbation responses in Perturb-seq are selected (columns). Expression changes (rows) are the union of top 20 differentially expressed genes (DEGs) for all perturbed genes.

To further enhance *in silico* screening performance, we extended the pertTF training strategy by incorporating prior biological knowledge through the lochNESS score (Fig. 6e). Specifically, known essential gene classes, such as ribosomal and proteasomal genes, were used to calibrate lochNESS predictions, enabling the model to further incorporate known biological knowledge into the training. Using this framework, pertTF accurately predicted strong negative lochNESS scores for essential genes. For example, MRPS5—a mitochondrial ribosomal protein not included in the training dataset—was predicted to have a strongly negative lochNESS score, consistent with its essential role in cellular viability (Fig. 6f). Across gene categories, predicted lochNESS scores showed clear separation between non-essential genes and essential gene classes, with ribosomal and proteasomal genes significantly enriched for negative lochNESS values (Fig. 6g). Finally, we evaluated pertTF’s ability to prioritize regulators of cell type composition in independent pooled CRISPR screens targeting definitive endoderm (DE) and posterior foregut (PFG, PP1) populations^39^. Genes whose knockout led to depletion or enrichment of the corresponding cell type in experimental screens were grouped as negatively or positively selected, respectively. Predicted lochNESS scores showed strong concordance with these experimental outcomes, with significantly lower scores for negatively selected genes and higher scores for positively selected genes in both DE and PFG screens (Supplementary Fig. 7b-e).

Finally, pertTF-generated predicted cell embeddings provide an approach to perform in silico Perturb-seq (Fig. 6h). We therefore generated in silico, single-cell perturbations on the genes whose perturbation induced strongest expression changes in our Perturb-seq dataset, and compared the predicted cell embeddings and expression change profiles with the actual perturbed cells in Perturb-seq (Fig. 6i-j, Supplementary Fig. 7f). pertTF generated cell embeddings are highly similar with true cell embeddings, where all tested genes are unseen perturbations from the training dataset (Fig. 6i). Perturbing genes from the same protein complex (e.g., SMARCC1 and SMARCD1) led to similar expression changes. In addition, cells with SALL4 and TCF7L1 perturbations cluster together, confirming the similar transcriptomic responses upon perturbing both genes^45^. For the expression prediction, pertTF predicted expression changes are highly similar with observed expression profiles (Fig. 6j; Supplementary Fig. 7f), including the ribosomal subunits whose expressions are up-regulated upon perturbing many chromatin regulators (e.g., SMAD4, SMARCA4, SMARCD1), consistent with previous observations in other Perturb-seq datasets^40^.

Together, these results demonstrate that pertTF enables accurate and biologically meaningful *in silico* genetic screens. By integrating embedding-based similarity and lochNESS-based composition prediction, pertTF can prioritize functional regulators, recover known biology, and generalize to genes do not present in the training dataset, providing a scalable computational alternative to experimental pooled CRISPR screening.

## Discussion

In this work, we present pertTF, a transformer-based framework for predicting cellular responses to genetic perturbations at single-cell resolution. By training pertTF on a uniquely curated perturbation-resolved scRNA-seq dataset spanning pancreatic differentiation, we demonstrate that the model can accurately predict not only gene expression changes, but also higher-order phenotypes such as cell type identity shifts and population composition changes. pertTF consistently outperforms existing foundation models and perturbation predictors across a wide range of tasks, including cell type classification, genotype inference, prediction in unseen cellular contexts, and generalization to unseen genes. Importantly, pertTF enables *in silico* genetic screening by ranking candidate gene perturbations according to their predicted phenotypic similarity or compositional impact, with strong concordance to independent pooled CRISPR screens. Together, these results establish pertTF as a generalizable and biologically grounded model for learning perturbation–response relationships in complex cellular systems.

A variety of computational strategies have been explored to predict single-cell transcriptomic responses to perturbation, each with distinct strengths and limitations^8–16^. Despite their promise, initial benchmarks have raised caution that current models do not yet decisively outperform simple baselines^29,30^ and have poor generalizability in unseen cellular contexts^31^. These findings underscore that capturing the full complexity of genetic perturbations and cellular context remains challenging. Our study represents one of the many ongoing efforts to generate high-quality perturbation data spanning different cellular and perturbation contexts (e.g., Tahoe 100M^46^, X-Atlas/Orion^47^, and many others^48,49^), improve model architectures and training strategies, or to incorporate additional biological priors to fully leverage the power of perturbation prediction models. Moreover, most models are still evaluated primarily on predicting expression changes in a limited number of systems (e.g., immortalized cell lines). pertTF is among one of the many ongoing efforts to predict higher-order phenotypic outcomes (e.g. multimodal readout^50^, changes in cell type proportions or functional phenotypes), and generalizability across different perturbations, cellular contexts and systems.

A key factor underlying the performance and robustness of pertTF is the quality and structure of the training and evaluation dataset. Many models and benchmark studies are trained on existing Perturb-seq datasets that focus on a single or limited number of cell types. In contrast, our dataset spans over 14 distinct cell types across multiple stages of pancreatic differentiation. This diversity enables pertTF to disentangle perturbation effects from intrinsic cell state variation and is critical for learning transferable perturbation representations. Equally important, the perturbations in this dataset are based on full gene knockouts, rather than variable and partial knockdowns, which are common for many CRISPRi-based Perturb-seq datasets. Complete loss-of-function perturbations in our dataset provide cleaner and more interpretable training signals, reducing ambiguity introduced by perturbation efficiency.

The unique perturbation dataset in this study also enables rigorous benchmarking paradigms for perturbation prediction models. We systematically evaluated pertTF under increasingly challenging conditions, including prediction in unseen cell types, unseen perturbation targets, and the joint setting of unseen genes in unseen cell types. Such benchmarks more closely reflect real-world use cases, where new perturbations or cellular contexts are often absent from training data. Furthermore, validation on independent experimental datasets, including primary human islet data and orthogonal perturbation modalities, provides strong evidence that pertTF predictions reflect genuine biological signals rather than dataset-specific artifacts.

While this study focuses on pancreatic development and beta-cell–relevant biology, the conceptual and methodological advances of pertTF are broadly applicable. Many disease-relevant cellular systems—such as primary human tissues, organoids, immune subsets, and neuronal populations—are experimentally challenging or prohibitively expensive to interrogate using large-scale perturbation screens. Limited cell numbers, ethical constraints, and technical barriers often preclude systematic experimental exploration of gene function in these settings. In this context, *in silico* perturbation models like pertTF provide an attractive and complementary alternative. By leveraging knowledge learned from well-characterized systems, such models can prioritize candidate genes, infer likely phenotypic outcomes, and guide targeted experimental validation. The ability of pertTF to generalize across cell types and perturbation targets, and to perform virtual genetic screens that align with pooled CRISPR results, underscores its potential utility as a hypothesis-generation engine across diverse biological and disease contexts.

Despite its strengths, this study has several limitations that point to important future directions. First, while the dataset is rich in cellular diversity, the number of perturbed genes remains relatively modest. Scaling pertTF to larger perturbation spaces will require larger and more comprehensive Perturb-seq datasets, ideally spanning hundreds to thousands of genes. Emerging technologies such as dCas9–MECP2-based repression, Cas12a multiplexed perturbations, and dual-guide CRISPR systems offer promising avenues to increase perturbation throughput while maintaining manageable experimental complexity. Second, although full knockouts provide clean training signals, many experimental and therapeutic settings involve partial or dosage-dependent perturbations. Extending pertTF to explicitly model perturbation strength and efficiency will be important for capturing more nuanced genotype–phenotype relationships. Incorporating perturbation modalities beyond CRISPR knockouts, such as CRISPRa/i, base editing, or pharmacological perturbations, is another natural extension. Finally, while pertTF integrates gene-level information through graph-based embeddings, future models may benefit from richer multimodal inputs, including chromatin accessibility, protein abundance, or spatial context. Such integrations could further enhance interpretability and predictive power, especially in systems where transcriptional responses alone are insufficient to explain phenotypic outcomes.

In summary, pertTF represents a step toward generalizable, context-aware AI models for genetic perturbation prediction. By combining high-quality perturbation data, rigorous evaluation, and biologically informed modeling, this work highlights both the promise and the challenges of using AI to build predictive, *in silico* representations of cellular systems.

## Methods

### pertTF method overview

#### model architecture

pertTF is a transformer-based neural network designed to model single-cell transcriptional responses to genetic perturbations. The transformer implementation of pertTF is largely inspired by scGPT^13^ and BERT. Briefly, the model takes as input a sequence of tokens (representing genes) for each cell, where each token is an encoding given by the combination of its gene identity (gene embeddings *e*^*g*^) and log normalized expression (value embedding *e*^*v*^). An extra empty token is appended to the beginning of the input token sequence to help store aggregate cell information and act as a cell embedding *e*^*c*^ in the transformer output, which serves as a shared backbone for downstream prediction tasks. An optional perturbation encoder, implemented as a learnable embedding layer, are added to the input of the model. The embedded tokens for each cell are then processed through multiple stacked transformer encoder layers, each consisting of a multi-head self-attention and feedforward networks. The core of pertTF employs a similar approach as previous models, through random masking of a portion of input tokens, pertTF optimizes a masked gene expression reconstruction loss *L*_*MSE*_ during training to encourage the model to learn biologically meaningful gene and cell representations that are informative for downstream tasks. This architecture enables pertTF to model high-order, non-linear gene–gene and gene–perturbation interactions while remaining agnostic to cell type–specific assumptions.

#### Key modifications

Compared with foundation models that also use transformer (e.g., BERT, scGPT, Geneformer, scFoundation), pertTF engineered several key modifications to capture multiple aspects of perturbation dynamics more effectively:

- **Perturbation integration:** A custom perturbation module directly integrates perturbation encodings into cell embeddings. The resulting **perturbed cell embeddings** are used to guide the accurate recreation of full gene expression profiles.
- **Distribution-aware modeling:** Instead of using mean squared error, cell-embedding-guided expression prediction is modeled using a negative binomial log-likelihood loss to better capture the underlying distribution of single-cell count data.
- **Biologically-targeted training:** During the masked gene expression reconstruction phase, pertTF samples tokens predominantly from pre-calculated highly variable genes (HVGs) at a 2:1 ratio against non-HVGs, prioritizing highly informative transcriptional signals.
- **Distinct latent spaces:** We implemented a supervised contrastive loss during classification. This forces the model to learn well-separated latent spaces for samples with different labels.
- **Adapters for phenotype/score prediction:** Adapters modules were added for prediction downstream predictions tasks such as cell type / genotype classification or prediction of composition changes.

#### Supervised contrastive learning for latent separability

To ensure the model learns highly separable latent representations for cell embeddings, we optionally incorporate a supervised contrastive loss. By leveraging available cell type and perturbation (genotype) labels as constraints, this objective encourages the clustering of cells with similar biological states while explicitly pushing apart embeddings from distinct cell types or perturbations. This forces the model to generate distinct, biologically meaningful clusters within the latent embedding space, improving the resolution of downstream tasks.

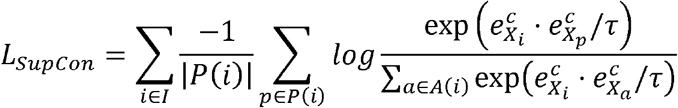

Where *P*(*i*) denotes the set of cells in a training batch with the same labels as cell *i*, and *A*(*i*) denotes all cells in the batch.

#### Classification Tasks

Classification of cell-type and genotype tasks were performed on the cell embeddings of each cell. Briefly, a fully connected neural network (FCNN) module was constructed for each classification task, and cell embeddings were used as an input to predict cell-type or genotype. Classification tasks were trained via a combination of the masked MSE, supervised contrastive loss and cross entropy loss:

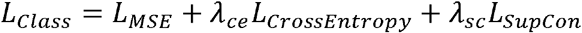

#### lochNESS score for cell type and composition changes

To quantify perturbation-induced shifts in cell identity or population composition, we adapted the lochNESS score [18] to evaluate each inquiry cell across all perturbations. For a specific perturbation and target cell type, the score is derived from the local neighborhood of wild-type (WT) and perturbed cells. We construct a *k*-nearest neighbor (kNN) graph, where distances to the inquiry cell are computed using the latent space embeddings. The lochNESS score and its background factor, σ_*bg*_, are computed as follows:

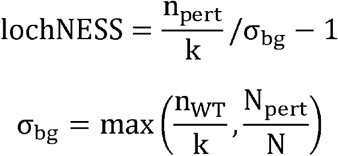

Where *k* is the total number of nearest neighbors; *n*_*pert*_ is the number of perturbed cells within the *k* -nearest neighbors; *n*_*wt*_ is the number of wild-type cells within the *k* -nearest neighbors; *N*_*pert*_ is the total number of perturbed cells in the dataset and *N* is the total number of cells in the dataset. The background factor is designed to avoid arbitrary large lochNESS scores due to either (1) there are very few numbers of perturbed cells in the whole dataset; or (2) the lochNESS score for wild-type cells are large for some cell types, which is counter-intuitive. Positive lochNESS values indicate enrichment of the perturbed cell, whereas negative values indicate depletion. lochNESS scores can be computed at the single-cell or population level and are used both as a prediction target during training and as a quantitative readout for downstream analyses, including virtual genetic screens. Prediction for the lochNESS score were optimized via mean-squared error (MSE) loss

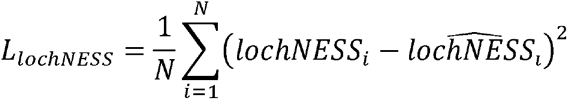

#### Gene expression prediction guided by cell embedding (GEPC) using negative binomial (NB) distribution

To better capture the overdispersed, count-based nature of single-cell RNA sequencing data, we adapted the scGPT architecture by replacing its standard masked mean squared error (MSE) objective with a Negative Binomial negative log-likelihood (NB-NLL) loss. In this formulation, the model decodes the distribution parameters for a given gene *A* within cell *X* Specifically, a neural network decoder processes the concatenation of cell *X*’s embedding and gene *A*’s embedding to predict an unscaled mean expression value. Concurrently, a gene-specific dispersion factor is decoded strictly from gene *A*’s embedding. The unscaled mean is then multiplied by the empirical size factor of cell *X* to establish the final mean of the Negative Binomial distribution. During training, the model minimizes the NB-NLL between these predicted parameters and the raw target transcript counts for each cell as shown:

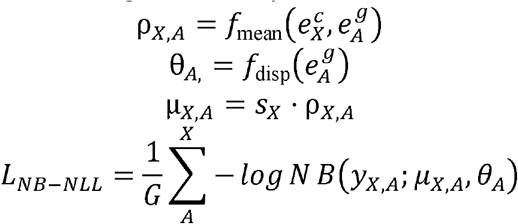

Where 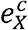 is the cell embedding, *e*_*A*_ is the gene embedding, *G* is gene sequence length, ρ is the unscaled mean, θ is the dispersion, *S*_*x*_ is the input cell size factor, μ is the final scaled mean, *y*_*X,A*_, is the raw count target and *f*_*mean*_ and *f*_*disp*_ are neural networks that predict the negative binomial parameters for gene *A* in cell *X*. For downstream inference and benchmarking, predicted counts are sampled directly from the parameterized Negative Binomial distribution and subsequently log-normalized.

#### Prediction of Perturbation Effects via pertTF

To enable the prediction of perturbation effects, pertTF utilizes a learnable embedding layer to encode genetic perturbations, denoted as *p*_*G*_ for a perturbed gene *G*. These perturbation embeddings—which can be *initialized initio* or derived from external representation models such as Graph Neural Networks (GNNs)—are concatenated with the non-perturbed (WT) cell embedding, 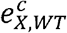, for a given cell *X*. This combined representation is then routed through a FCNN module to generate a corresponding perturbed cell embedding, 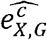 .

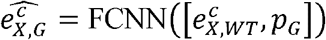

Finally, this perturbed cell embedding is passed to the GEPC module to reconstruct the predicted post-perturbation gene expression profile. During training, each non-perturbed cell is randomly paired with a perturbed cell of the same celltype. The loss for perturbation expression prediction is as follows:

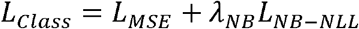

#### Predicting perturbation effects in new cellular contexts

During training, the model has access to all cells of the training set as well as WT cells of the held-out cell types, meaning only perturbed cells of the held-out cell types are unseen. Post training, the WT cells of the held-out cell types were perturbed accordingly to compare with the true perturbed cells, thus evaluating the model’s ability to generalize perturbation effects across cellular contexts rather than memorizing cell type–specific responses.

#### Predicting unseen perturbations

To predict perturbation effects for genes that are not present in the training dataset, pertTF incorporates external gene embeddings derived from a graph neural network (GNN), implemented in the published GEARS^6^ algorithm. Briefly, the GNN is trained on a gene–gene interaction graph constructed from curated Gene Ontology (GO) databases. Each gene is represented as a node, and edges encode functional or regulatory relationships. The resulting GNN embeddings capture functional similarity among genes and are used to parameterize perturbation embeddings for unseen genes. The perturbation GNN embeddings are first trained on an unrelated, published K562 Perturb-seq dataset^21^. These embeddings are then passed to pertTF as frozen values for perturbation embeddings, and trained together with other components of pertTF to generate predicted perturbed cell states, enabling extrapolation to novel perturbations.

#### Training setup

All training for pertTF models were done using NVIDIA H200 machines. Flash attention was utilized to accelerate the training process and accommodate a batch size of 128 for larger models. All models were optimized via Adam with default parameters, and an epoch-wise step schedular with learning rate decay at 0.995. Additionally, instead of randomly selecting a subset of genes for each cell every epoch during training, we pre-calculated 5000 highly variable genes (HVGs) on the training data and predominantly sampled genes from the HVGs set during training (ratio of HVG to non-HVG is set as 2:1, i.e. for a context length of 3000 tokens, 2000 are HVGs and 1000 are non-HVGs), to make sure every cell’s token sequence contained predominantly HVGs along with a smaller portion of non-HVGs. 5000 HVGs are recalculated separately for all validation data and are used in their entirety (along with a random sample of 1000 non-HVGs to construct a length 6000 token sequence) during evaluation. We found that this setup significantly improved the performance of downstream tasks.

### Training and validation data generation

#### Training data

Training data is generated by NIH MorPhiC (Molecular Phenotypes of Null Alleles in Cells) consortium and is available at the MorPhiC data portal (https://morphic.bio/). Data were generated using pooled CRISPR-based gene knockout experiments in human pluripotent stem cells (hPSCs) undergoing directed pancreatic differentiation. A total of 30 genes, including developmental regulators and diabetes-associated factors, were targeted using loss-of-function perturbations. Successfully transduced cells were enriched by fluorescent sorting and differentiated in pooled cultures. Single-cell RNA sequencing was performed at multiple differentiation stages, capturing over 87,000 high-quality cells across 14 annotated cell types. Perturbation identities were confirmed using targeted genotyping and guide barcode demultiplexing. Only cells with high-confidence perturbation assignments were included for training.

### Perturb-seq (50-gene)

DNA oligonucleotide pools (Twist Bioscience) were cloned into the LentiGuide(10X)-BFP-Puro backbone modified with a 10x Genomics Capture Sequence at the 3’ end and electroporated into Endura Duo competent cells (Lucigen) to generate pooled sgRNA libraries. For lentiviral production, 8 µg of sgRNA library plasmid was co-transfected into HEK293T cells with 6 µg psPAX2 and 2 µg pMD2.G packaging plasmids (Addgene #12260 and #12259, respectively). Media was replaced the following day, and viral supernatant was harvested 48 hours later^51^.

For viral transduction, WTC11-CLYBL-dCas9-KRAB cells were dissociated into single cells and infected in suspension at ∼80% confluency in the presence of CloneR2. Following recovery and expansion, BFP-positive cells were enriched by fluorescence-activated cell sorting 8 days post-infection, expanded, and cryopreserved in liquid nitrogen.

Frozen cells were rapidly thawed in a 37 °C water bath, pelleted, and washed 2–3 times with PBS supplemented with 2% FBS and 5 mM EDTA (1 mL per sample). Cells were resuspended in PBS containing 2% FBS and 5 mM EDTA (200 μL per 1–1.5 × 10□ cells) and stained with LIVE/DEAD™ Violet Viability/Vitality Kit (L34958; Invitrogen) at a 1:1000 dilution for 10 min on ice. After staining, cells were resuspended in PBS with 2% FBS and 5 mM EDTA (500 μL per 1–1.5 × 10□ cells).

Live single cells were isolated by fluorescence-activated cell sorting using a BD FACSymphony™ S6 Cell Sorter (BD Biosciences), followed by one wash with PBS supplemented with 0.04% BSA (1mL per sample). Single-cell RNA-seq and sgRNA capture libraries were generated using the Chromium Next GEM Single Cell 3′ HT Reagent Kit v3.1 (Dual Index; 10x Genomics) according to the manufacturer’s instructions. We prepared two biological replicates. For each replicate, approximately 110,000 cells were loaded, split across two Chromium chip lanes (10x Genomics). Two libraries were prepared from each Chromium lane: a 3’ gene expression library for single cell RNA-seq and a CRISPR screening library for sgRNA capture. All the libraries were sequenced on an Illumina NovaSeq 6000 platform (Illumina). Each 3’ gene expression library was sequenced to reach ∼20,000 read pairs per cell, and each CRISPR screening library for sgRNA capture was sequenced ∼50-60 × 10□ reads.

### Classification Benchmarking

PertTF’s celltype and genotype classification performance was benchmarked against scGPT, Geneformer and scFoundation. 5-fold cross validation was performed for all methods on the full training data. Macro F1 score, macro average precision (PR-AUC) and Accuracy (ACC) metrics were used to evaluate classification performance. A separate model for all benchmarked methods was trained using the full training data and used to evaluate performance on the independent validation data.

### Perturbation Prediction Benchmarking

pertTF was benchmarked against state-of-the-art models for perturbation prediction, including scGPT, GEARS and scFoundation. All models were evaluated using identical training and test splits where applicable.

Specifically, when benchmarking against scGPT and GEARS, all training and prediction was performed for the top 5000 HVGs along with perturbed genes in the training data and independent datasets. Benchmarking against scFoundation was performed for ∼20,000 genes due to scFoundation’s model requirements.

Benchmarks were performed under two settings. In the first, the training data was split into 5 distinct sets, including a training set of 80% of perturbations and 80% of cell types, a validation set of 5 % of perturbations, and unseen cell types (20% of cell types), unseen perturbations (15% of perturbations), and a joint set of unseen perturbations and unseen cell types. In the second setting, all the training data was used during training and used to evaluate the independent data.

The cell-eval framework was used to facilitate the evaluation of all methods. Of the cell-eval metrics, we selected 8 informative metrics, as described in [STATE paper], based on prevalence in benchmarking studies/contests and methodology literatures. Briefly, these included overlap of all differentially expressed genes (Overlap at N), differential expression direction matching (DE-direction), ROC-AUC, PR-AUC, Pearson Correlation Coefficient of expression changes (PCC-delta), mean absolute error of expression (MAE), MAE of expression change (MAE-delta) and L1 Discrimination Score (additional details in Supplementary). These metrics were calculated independently for each perturbation of each annotated celltype on the predicted vs ground truth cells. Moreover, we include a cosine similarity metric to evaluate cell embeddings of predicted and true perturbed cell.

### Application to primary human islet data

Two published primary human islet datasets were used^19,20^. In the first dataset^19^, primary human islet scRNA-seq datasets from non-diabetic, pre–Type 2 Diabetes, and T2D donors were downloaded from the Gene Expression Omnibus (GEO) accession number GSE200044. Only the scRNA-seq part of the multiome datasets were used. A pre-trained pertTF model was fine-tuned using a small number of labeled primary cells for cell type annotation. Following fine-tuning, pertTF was applied to infer latent perturbation states, predict perturbation probabilities, and compute lochNESS scores across donors and cell types. In the second dataset^20^, processed, scMultiome data was downloaded from the corresponding Zenodo repository (https://zenodo.org/doi/10.5281/zenodo.6515986). Predicted perturbation states were validated by examining concordance with known transcriptional signatures and independent perturbation experiments, including siRNA-mediated knockdowns.

### Virtual genetic screening

Virtual genetic screens were performed using two complementary strategies. In the first approach (Method 1), predicted cell embeddings for candidate perturbed genes were compared with a target perturbed population using cosine similarity, and genes were ranked from the highest similarity to the lowest similarity. In the second approach (Method 2), genes were ranked using predicted lochNESS scores to directly estimate enrichment or depletion of target cell populations. These rankings were validated against two sources: (1) known pan-essential genes and non-essential genes, defined in ref ^25^ and MAGeCKFlute^26^; (2) published pooled CRISPR screens to identify regulators during pancreatic differentiation^24^, including definite endoderm (DE) screens focusing on the expression of SOX17-GFP (a DE marker) and pancreatic progenitors (PP) screen, targeting the expression of PDX1-GFP (a PP marker). To improve robustness, known essential gene categories, such as ribosomal and proteasomal genes, were incorporated during training to calibrate lochNESS score distributions, by setting up the corresponding lochNESS score to -1 for essential genes (to simulate depletion effect of cell composition). In contrast, known non-essential genes are also incorporated, by setting the corresponding lochNESS score to 0 (i.e., unchanged cell composition).

## Acknowledgements

We would like to thank all members of Li laboratory and Huangfu laboratory, and NIH IGVF and MorPhiC consortiums for comments and discussions. W.L. is supported by funds through NIH (R01HG010753, R01HL168174), the National Cancer Institute—Cancer Center Support Grant (CCSG) – P30CA134274, and startup fund from University of Maryland Institute for Genome Sciences (IGS), Pharmacology and Physiology, and Medicine Institute for Neuroscience Discovery (UM-MIND). W.L. is an investigator at the University of Maryland– Institute for Health Computing (UM-IHC), which is supported by funding from Montgomery County, Maryland, and the University of Maryland Strategic Partnership: MPowering the State, a formal collaboration between the University of Maryland, College Park and the University of Maryland, Baltimore. D.H. is supported by grants from NIH (UM1HG012654, U01HG012051, R01HD111256) and MSKCC Cancer Center Support Grant from NIH (P30CA008748).

## Data availability

Knockout village scRNA-seq raw sequencing data are available in ERP168459 and the processed read counts are available at the GEO under accession code GSE313516. The processed Seurat object and metadata including cell type and genotype annotation is available on Zenodo (https://doi.org/10.5281/zenodo.17824935). Datasets are also available in Hugging Face website: https://huggingface.co/weililab.

## Code availability

The source code of pertTF is available, open source, under the MIT license, at https://github.com/davidliwei/pertTF. The trained model is available at the Hugging Face website: https://huggingface.co/weililab.

## Supplementary Figures

**Supplementary Figure 1**. Training pertTF. **a-b**. The loss of metrics in training data (a) and validation data(b) including cell type classification (cls), perturbation classification (pert), and masked expression, measured by mean squared error (mse) and mean absolute error (mre). **c**. The predicted cell type and genotype of each cell, corresponding to the actual cell type and genotype in Figure 1e-f. **d-e**. The Area Under the Receiver-Operating Characteristic Curve (AUC) and Area Under the Precision-Recall Curve (AUPR) for cell type classification (d) and perturbation classification (e).

**Supplementary Figure 2**. pertTF cell embeddings after incorporating perturbation information as input. **a-b**. Cell embeddings from typical scRNA-seq pipeline using top 2000 highly variable genes, followed by PCA and UMAP clustering. **c-d**. Cell embedding clustering without masking perturbation information. **e**. pertTF generated cell embeddings by varying the weights of loss function in perturbation classification (cross entropy). Only data from day 18 (late-stage differentiation cell types) are used to evaluate the impact of perturbation weight. **f-g**. pertTF generated cell embeddings on all cell types.

**Supplementary Figure 3**. Cell composition changes. **a**. The lochNESS score distribution of GATA4 and HHEX across all major cell types. **b**. The predicted and true TADA2B lochNESS score. **c**. A comparison of lochNESS score prediction using pertTF and a naïve gene expression within different genotypes. **d**. Cell embeddings of pertTF in Figure 2f, colored by cell types. **e**. Performance comparisons between pertTF and scFoundation. Since the input dimension of scFoundation (∼20,000) is different from all other methods, the comparison is performed separately from Figure 2g.

**Supplementary Figure 4**. Performance comparison on unseen perturbations. **a**. The assigned cell types of PDX1 perturbation prediction, corresponding to Figure 3b. **b**. A comparison of predicted vs true cell embeddings of different genotypes using pertTF and scGPT. **c**. A comparison of pertTF and scFoundation on metrics of gene expressions for unseen perturbations. **d**. A comparison of pertTF and scFoundation on metrics of gene expressions for both unseen cell types and perturbations.

**Supplementary Figure 5**. Independent experimental validations using Perturb-seq. **a**. A UMAP presentation of all cells in Perturb-seq. **b**. Mixscape classified cells for CTNNB1 perturbed cells into “NP” (non-perturbed) or “KO”. Both “NP” and “KO” are classification labels generated by Mixscape. **c**. The calculated PS score of CTNNB1 perturbed cells. **d**. A comparison of pertTF and scFoundation. **e**. The predicted cell embeddings of CTNNB1 from scGPT. **f**. A comparison of cosine similarity values between predicted vs true cell embeddings of CTNNB1 using different methods.

**Supplementary Figure 6**. Application of pertTF to primary cells. **a**. The percentage of cells classified as wild-type or gene loss across different individuals with non-diabetes, pre-T2D and T2D. **b**. The expression of signature genes in NEUROD1 knockouts vs wild types in SC-beta cells in the training dataset. **c**. The signature gene expressions of HNF4A and RFX6 across cells with different groups of probabilities that have HNF4A or RFX6 perturbation (P_HNF4A_ or P_RFX6_).

**Supplementary Figure 7**. In silico screen for DE and PP screens. a. The precision-recall curve for pertTF vs. gene expression only. The number indicates the Area Under the Precision-Recall Curve (AUPR) value for each method. **b-c**. The cumulative distribution of predicted lochNESS scores on the top hits of PP screen. Genes whose knockout deplete (or enrich) the corresponding DE/PP cell type in pooled screen are grouped as neg. (or pos.) selected, respectively. **d-e**. The distribution of predicted lochNESS scores on the top hits of DE screen. **f**. The predicted vs true expression changes of perturbing SMARCA4 at a single-cell level.

## Notes

https://github.com/davidliwei/pertTF

https://huggingface.co/weililab

### Competing Interest Statement

The authors have declared no competing interest.

