## Supplementary figures and images for "pertTF: context-aware AI modeling for genome-scale and cross-system perturbation prediction"

### Supplementary Figures

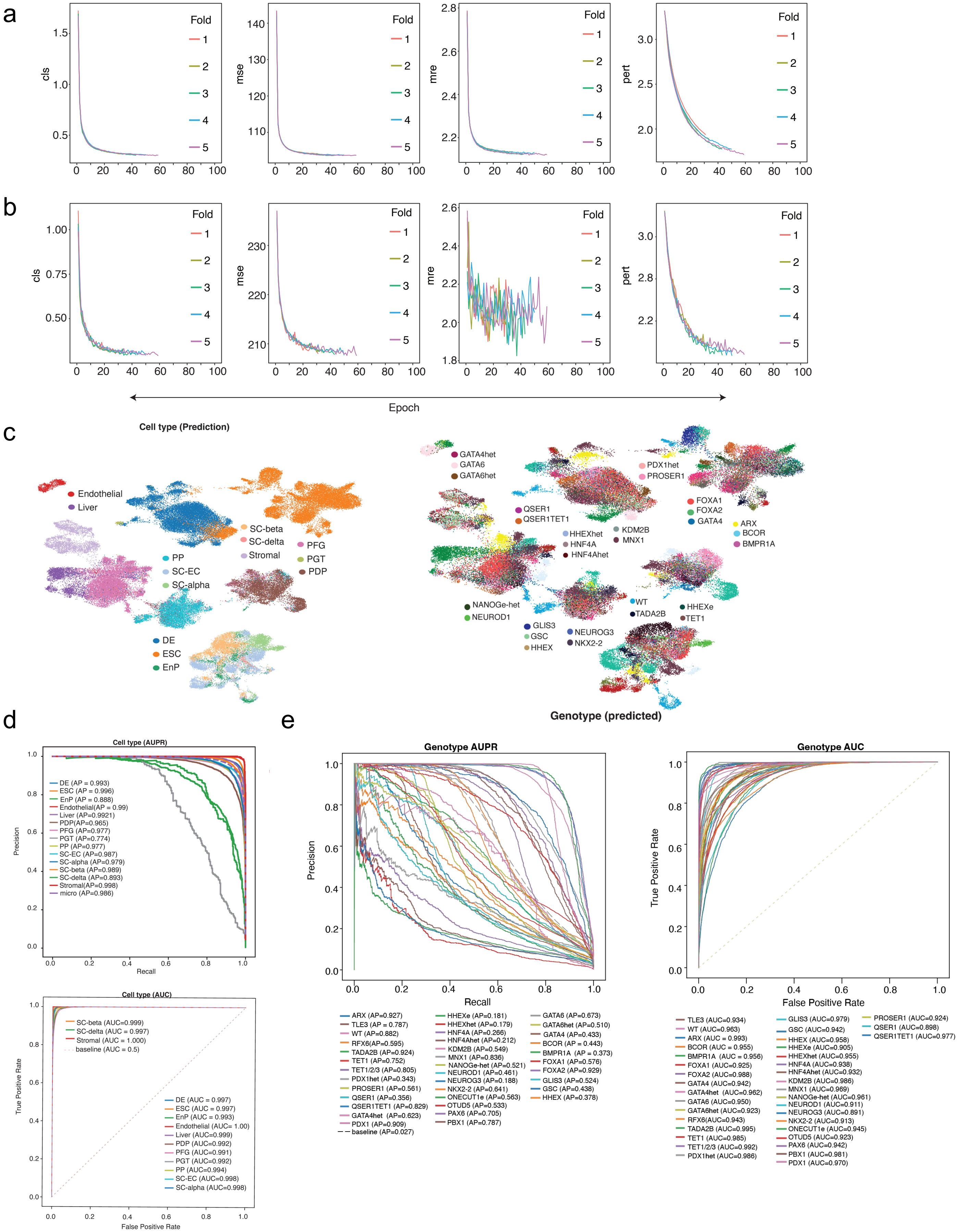

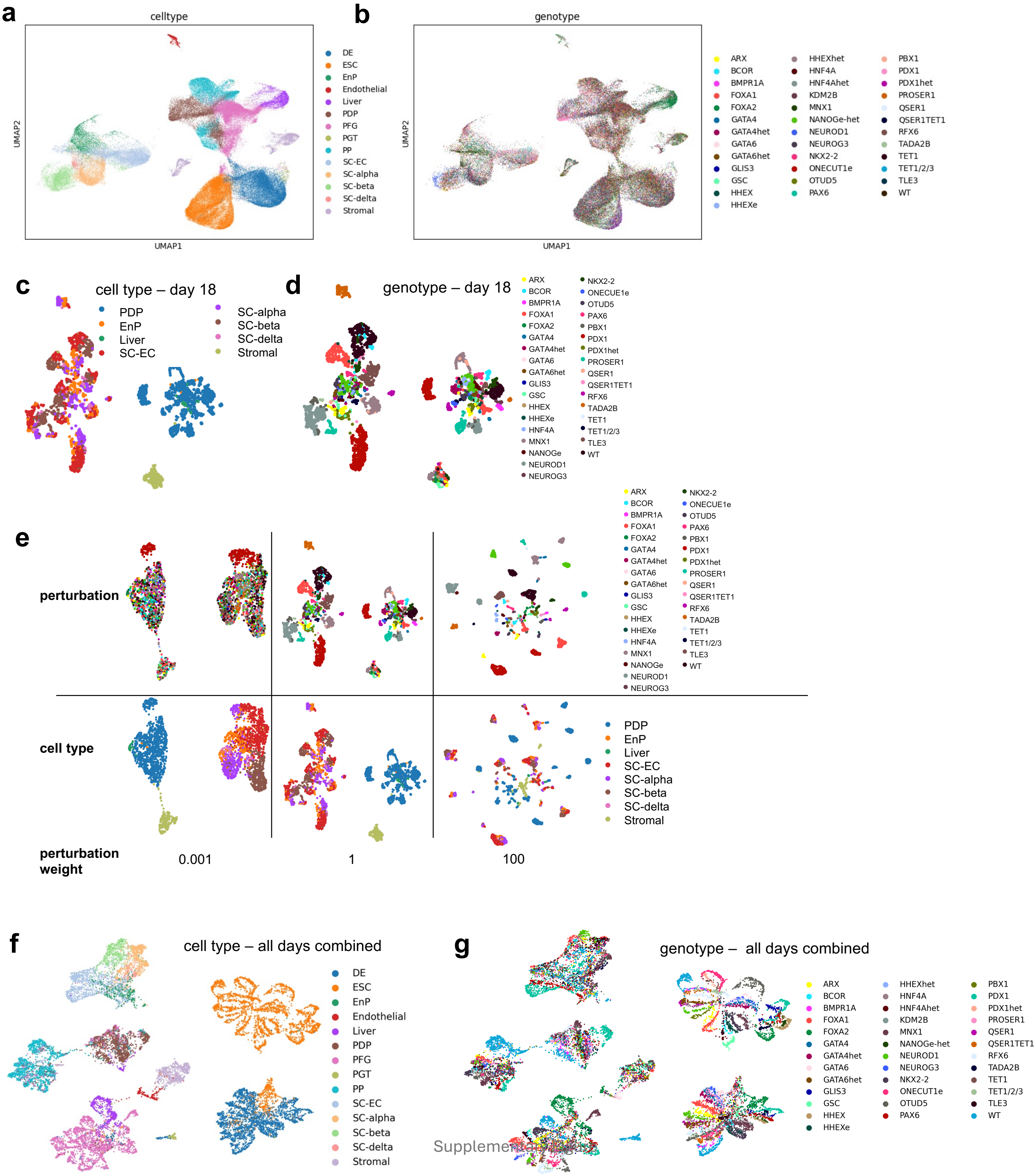


Supplementary Figure [illegible]

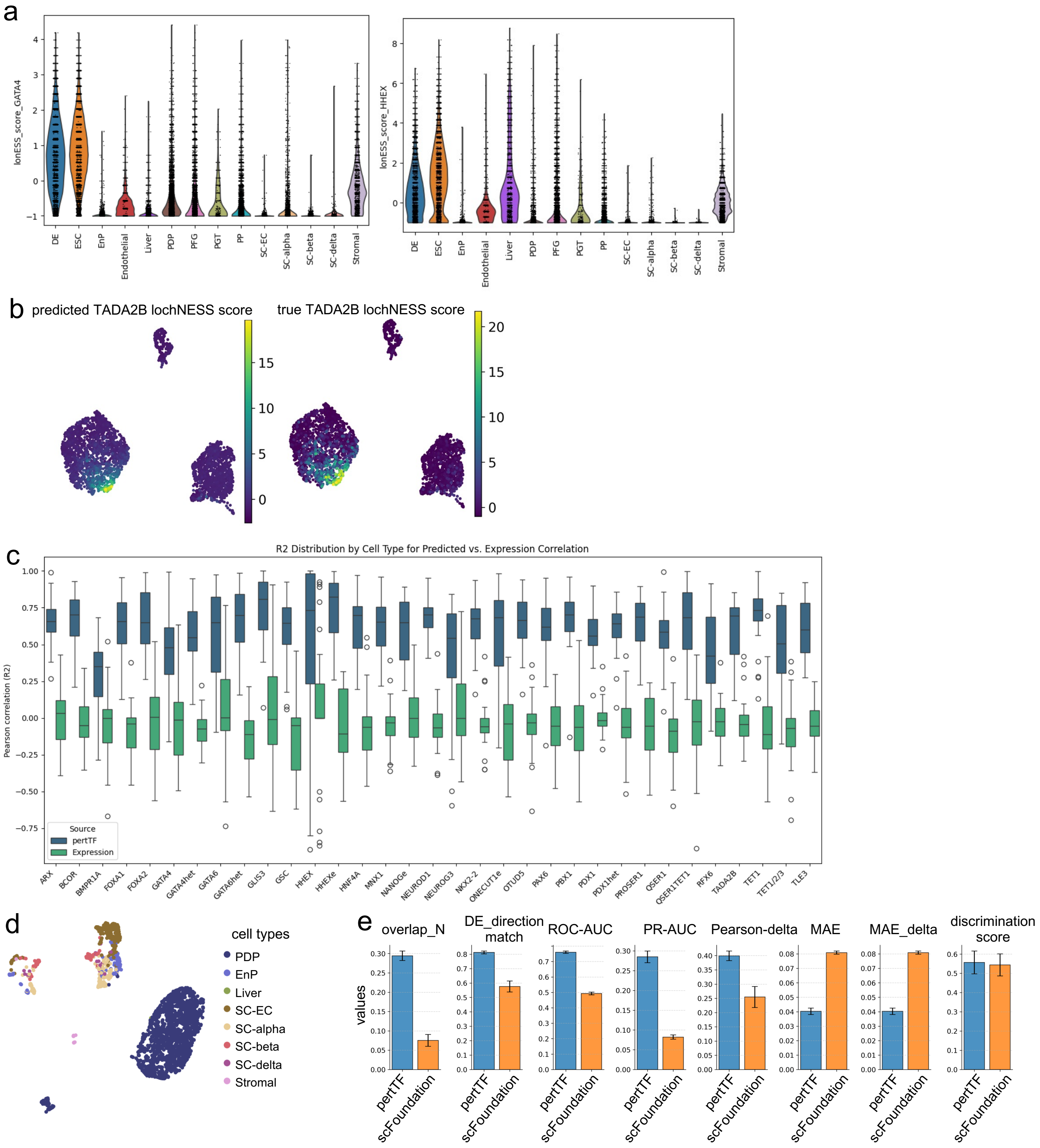

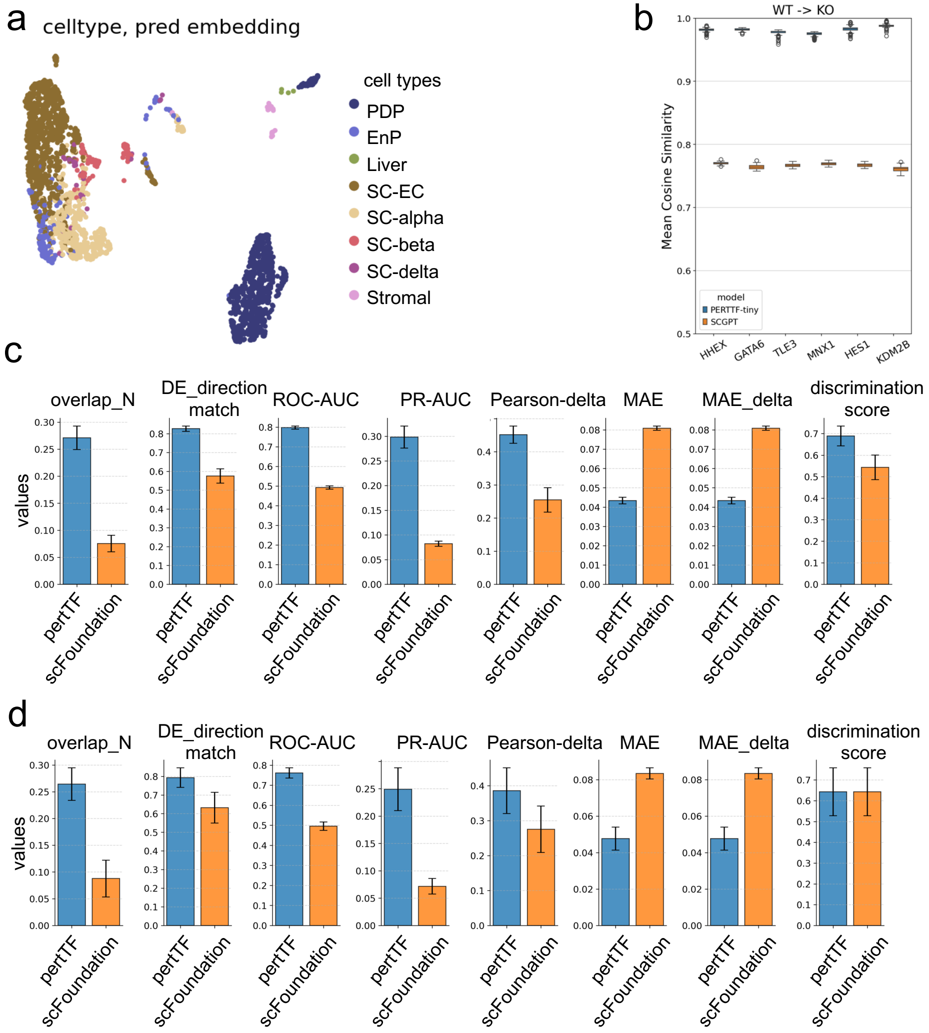

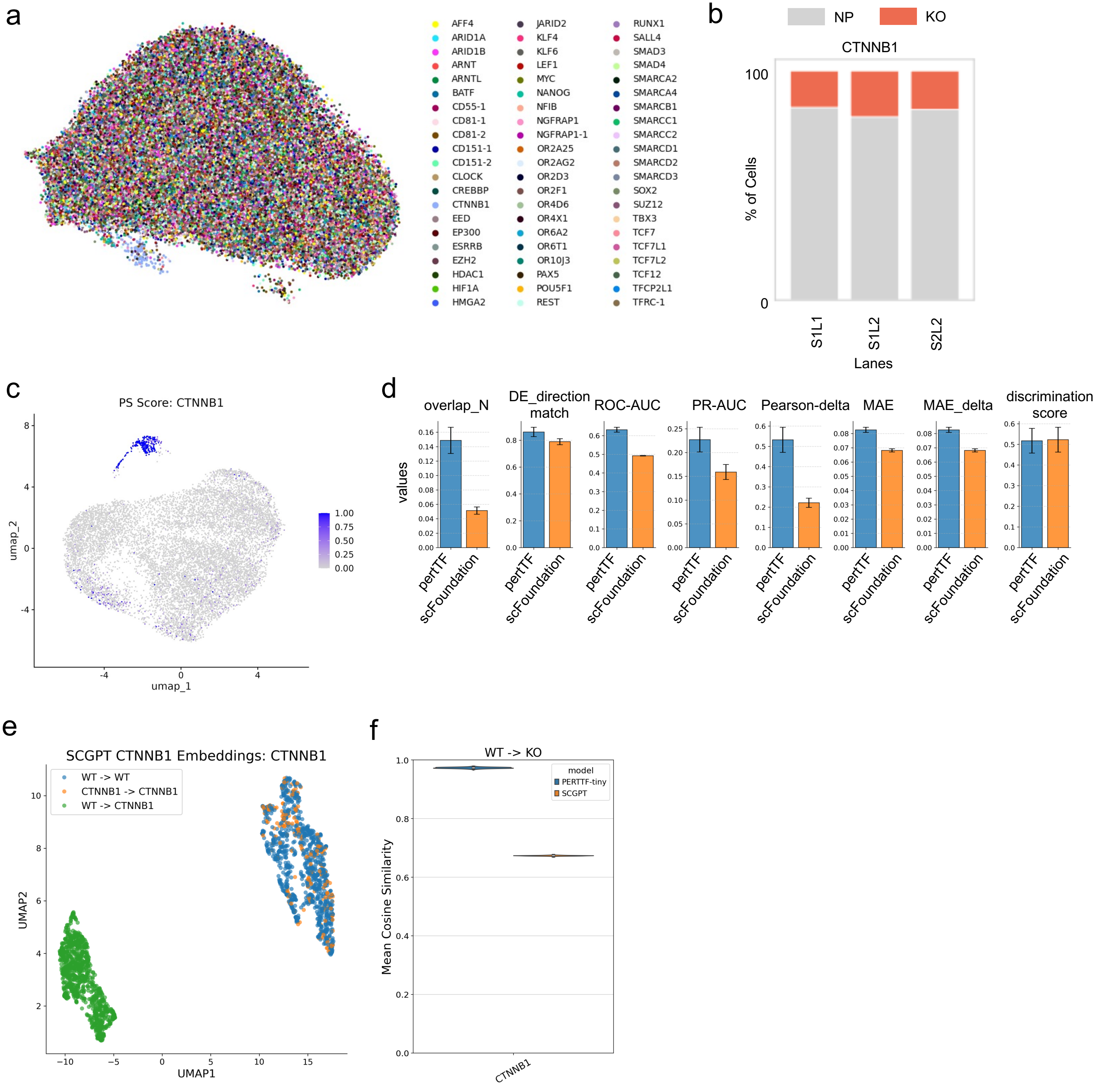

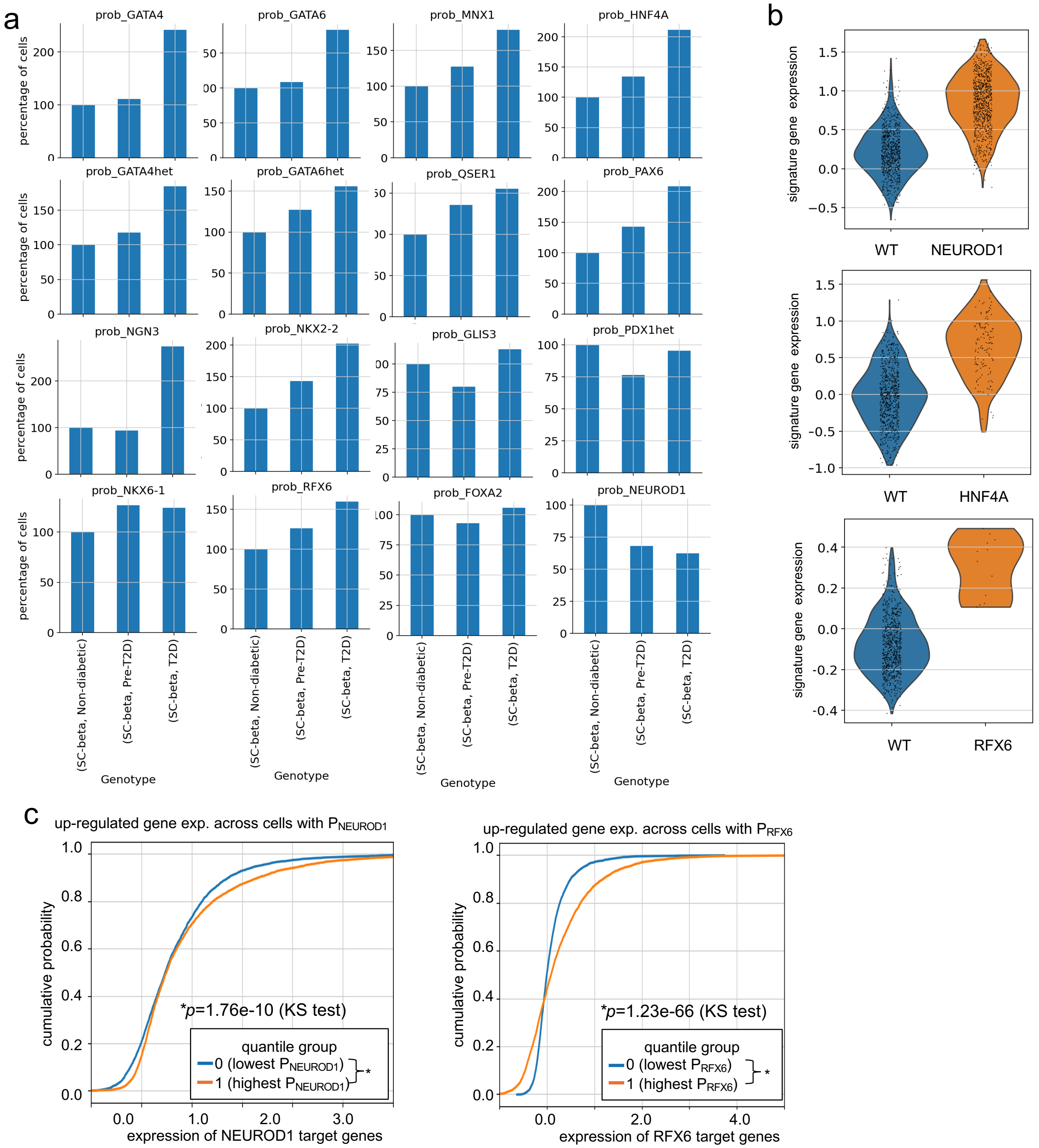

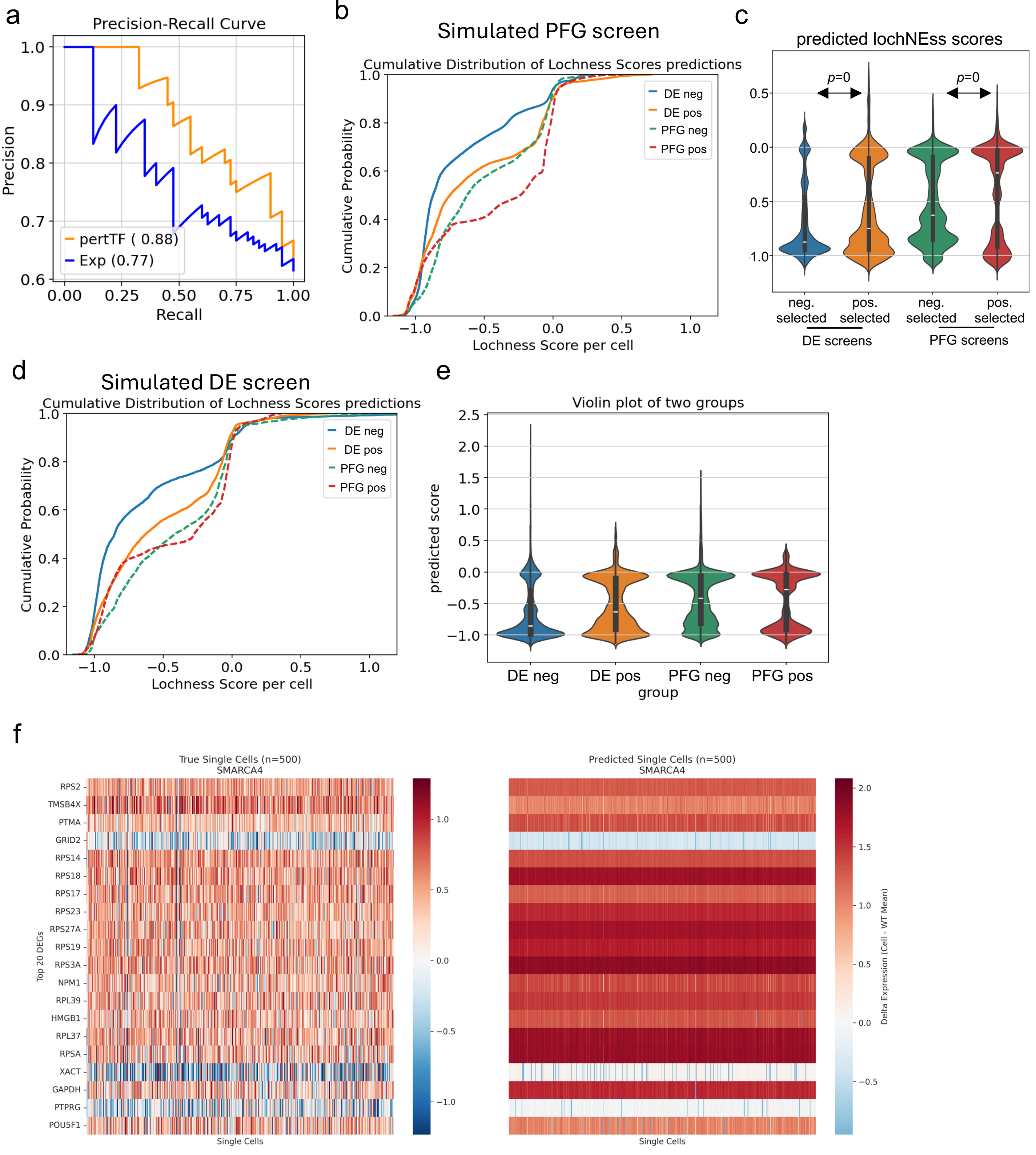
